# Maternal preconception calorie restriction reprograms coping strategies, socio-sexual behaviour, and endocrine function in adult rat offspring

**DOI:** 10.64898/2026.02.25.706709

**Authors:** Matthew D Zelko, Agnes Hazi, Helen Nasser, Elizabeth A Levay, Michelle Corrone, Jim Penman, Terrance G Johns, Antonina Govic

## Abstract

Maternal nutrition before conception is recognised as a determinant of offspring development; however, the behavioural and neuroendocrine consequences of preconception calorie restriction (CR) remain poorly understood. This study isolated the preconception window to examine how different CR patterns, stable (25% reduction; CR-25%), unpredictable deprivation (CR-A), and variable (25-75% fluctuation; CR-V), affect adult offspring outcomes. Male and female progeny from preconception CR female Wistar rats were assessed across domains sensitive to early-life programming, including anxiety- and depression-like behaviour, coping style, socio-sexual behaviour, and hypothalamic-pituitary-gonadal (HPG) axis activity. Preconception CR produced sex- and diet-specific effects. Females exhibited transient reductions in exploratory behaviour and more active coping styles, particularly CR-25% and CR-V animals. In males, all CR regimens enhanced copulatory behaviour and reduced aggression toward females. Endocrine profiling revealed divergent HPG responses: CR-A males showed elevated basal faecal testosterone metabolites (fTM) but reduced basal serum testosterone, whereas CR-V males exhibited blunted androgenic reactivity post-social provocation. These findings demonstrate that maternal preconception CR can program male offspring toward a prosocial, sexually motivated phenotype and female offspring toward an enhanced coping style, underscoring this period as a sensitive window for shaping behavioural and endocrine trajectories.

## 1 Introduction

It is well established that caloric and nutrient imbalances during pre- and postnatal development are potent environmental stimuli capable of exerting profound and lasting effects on offspring physiology and behaviour [1–8]. Emerging evidence suggests that the preconception period also represents a critical window in developmental programming, during which maternal nutritional status can influence oocyte maturation, gamete quality, the uterine environment, and epigenetic modifications that shape the trajectory of offspring development [9,10]. Indeed, nutritional perturbations occurring even before conception are increasingly recognised as contributors to adverse long-term health outcomes in progeny [11].

Research investigating maternal preconception nutritional status has largely focused on micronutrient deficiencies, such as folate [12] and iron [13], macronutrient deficiencies, such as protein restriction [14,15], and maternal adiposity [16]. These studies demonstrate that maternal nutritional deficiencies prior to conception can influence metabolic outcomes [15], fat deposition [17], neurodevelopment [12], and the emergence of neuropsychiatric disorders in adult progeny [16]. However, while these studies provide compelling evidence for preconception programming effects, few studies explicitly isolate the preconception period from the broader perinatal developmental window, and even fewer examine the impacts of global calorie restriction (CR) per se [6,18].

Another relatively unexplored area of the preconception-sensitive window is how global CR shapes offspring behaviour and neuroendocrine outcomes. While some studies have implemented maternal food restriction prior to conception, these diets often continue through pregnancy and/or lactation [19,20], making it difficult to disentangle the unique effects of the preconception period from those of the perinatal environment. Nevertheless, limited evidence indicates that even brief maternal preconception food restriction can influence offspring behavioural outcomes. For instance, a three-day maternal CR administered approximately one week before mating increased anxiety-like and risk assessment behaviour in adult male offspring in the open field and emergence tests [18], while also enhancing their copulatory efficiency [6].

In addition to the limited isolation of the preconception period, there has been minimal investigation into how different patterns of restriction – reflecting real-world variability in food access – might differentially program offspring behavioural outcomes. In such scenarios, maternal nutritional status may be consistently limited (stable CR), include intermittent episodes of acute food shortages (unpredictable deprivation), or fluctuate unpredictably (variable CR), each potentially exerting distinct effects on developmental programming. To address these differential patterns, three preconception CR regimens were implemented: a stable 25% reduction, unpredictable intermittent episodes of deprivation lasting 6 - 48 hours, and a variable restriction alternating between mild and moderate reductions (25-75%). Constant CR25% allowed us to examine the effects of a stable, moderate energy deficit. The repeated food deprivation group modelled episodic starvation, and the variable CR group introduced unpredictability in the maternal environment, mimicking real-world nutritional instability.

Collectively, this study aimed to address several outstanding questions regarding maternal preconception nutritional programming by isolating the preconception-sensitive window, systematically varying dietary restriction patterns, and examining behavioural outcomes in adult offspring. Following these preconception manipulations, offspring were assessed across several domains, including anxiety- and depression-like behaviour, coping style, socio-sexual behaviour, and concentrations of testosterone and faecal testosterone metabolites (fTM): outcomes known to be sensitive to early-life stress and maternal metabolic and dietary signals [21,22]. Given the modulatory role of postnatal maternal care on the behavioural profile of offspring [23] and the capacity for maternal nutritional disturbances to modify caregiving [18], maternal behaviour was monitored to better contextualise offspring outcomes. Outcomes were examined in both male and female offspring, as parental perturbations during the preconception period, including stress [24] and dietary alterations [25], can have sex-specific effects. Clarifying how maternal preconception dietary restriction shapes offspring neurobehavioural outcomes will advance our understanding of the early-life origins of risk and resilience and inform future preventative strategies.

## 2 Results

### 2.1 Body Weight

#### 2.1.1 Body Weight: F0 Females

The body weight of F0 females was measured periodically across five phases: acclimation, diet, mating, gestation and postnatal/lactation period. Figure 1 shows weights for each group by phase, where control females exhibited a steady increase in body weight from acclimation through to gestation, with a more pronounced rate of weight gain during gestation. A marked reduction in body weight occurred at parturition, followed by a gradual increase during the postnatal phase. All dietary interventions followed a similar trajectory; however, ROPE comparisons from a linear growth model indicated that both CR-25%, [E_m_ = 1.06 [0.72, 1.42], D_p_ = 100, ROPE_p_ = 0] and CR-V [E_m_ = 1.57 [1.23, 1.91], D_p_ = 100, ROPE_p_ = 0] exhibited a meaningful reduction in body weight growth compared to control F0 females during the dietary intervention (“Diet” phase).

**Figure 1.**
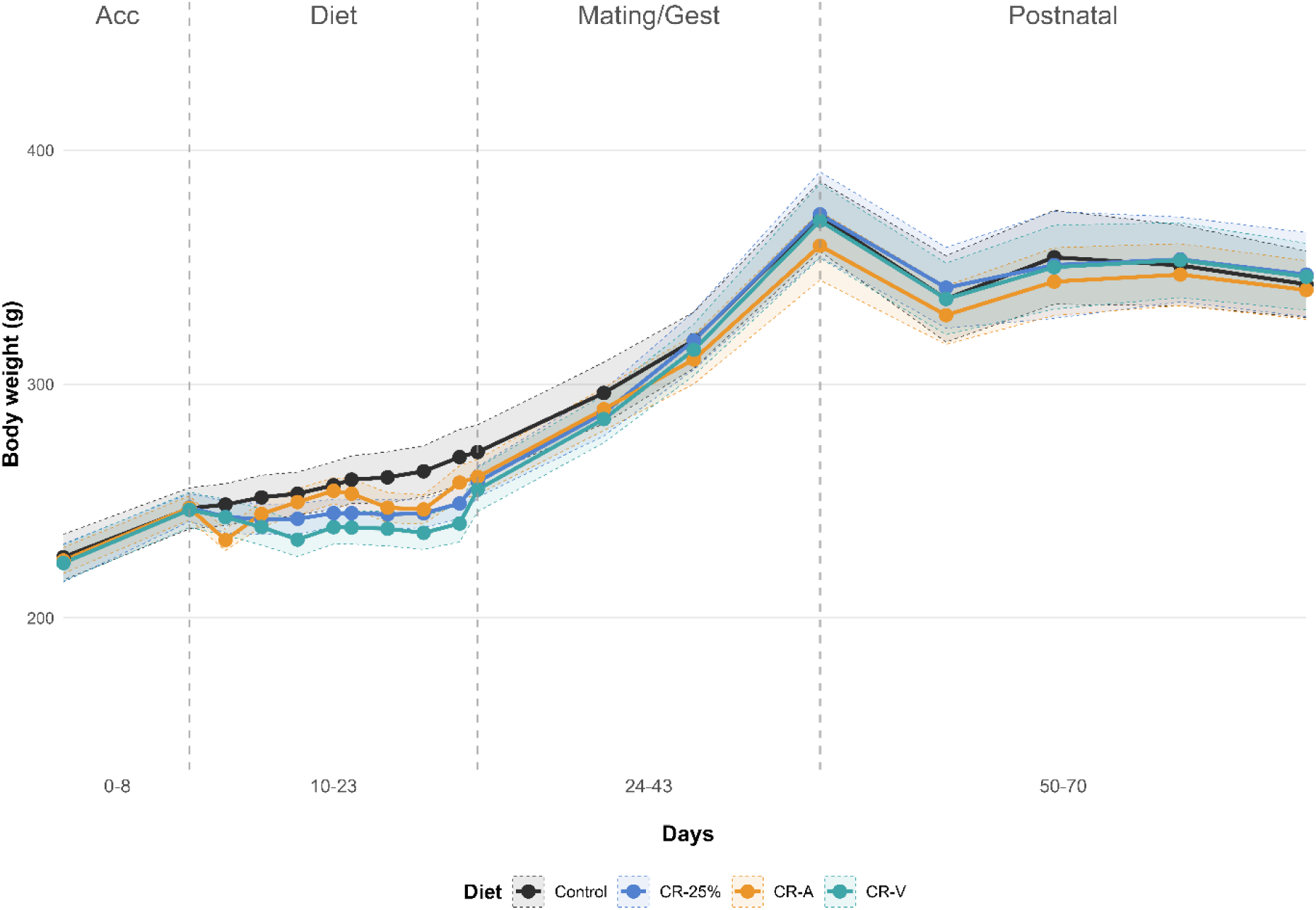
F0 Female body weight across acclimation, dietary restriction, mating, gestation, and postnatal phases. Solid lines indicate mean body weight trajectories for each diet, and shaded regions with dashed edges represent the 95% confidence interval of the mean. Vertical dashed lines indicate transitions between the Acclimation (Acc), Dietary Restriction (Diet), Mating/Gestation (Mating/Gest), and Postnatal periods. Note: Time period windows are to scale, but days (e.g., 36-43) indicate when first and last measurements for that period were taken.

#### 2.1.2 Body Weight: Offspring

Offspring body weight for the first 21 PND increased linearly across all maternal diet groups, regardless of sex, and in line with the growth of the control diet (see Figure 2A below). Growth continued throughout PND 32 to 129 in both sexes, with all cohorts demonstrating similar increases in weight over time. Figure 2B below visualises this growth and similarity for cohort 3 as an example of the time series, and ROPE comparisons confirmed that maternal diet had no meaningful effects on offspring body weight during the first 129 days. Thus, maternal preconception diet did not influence post-weaning growth in either sex. ROPE comparisons by cohort did confirm that, on average, males weighed more than females from PND 32 onwards, with the log median difference for cohort 1 [E_m_ = 0.31 [0.27, 0.36], D_p_ = 100, ROPE_p_ = 0], cohort 2 [E_m_ = 0.36 [0.32, 0.39], D_p_ = 100, ROPE_p_ = 0], and cohort 3 [E_m_ = 0.45 [0.42, 0.48], D_p_ = 100, ROPE_p_ = 0. This indicates that maternal diet did not affect the expected sex differences in weight.

**Figure 2.**
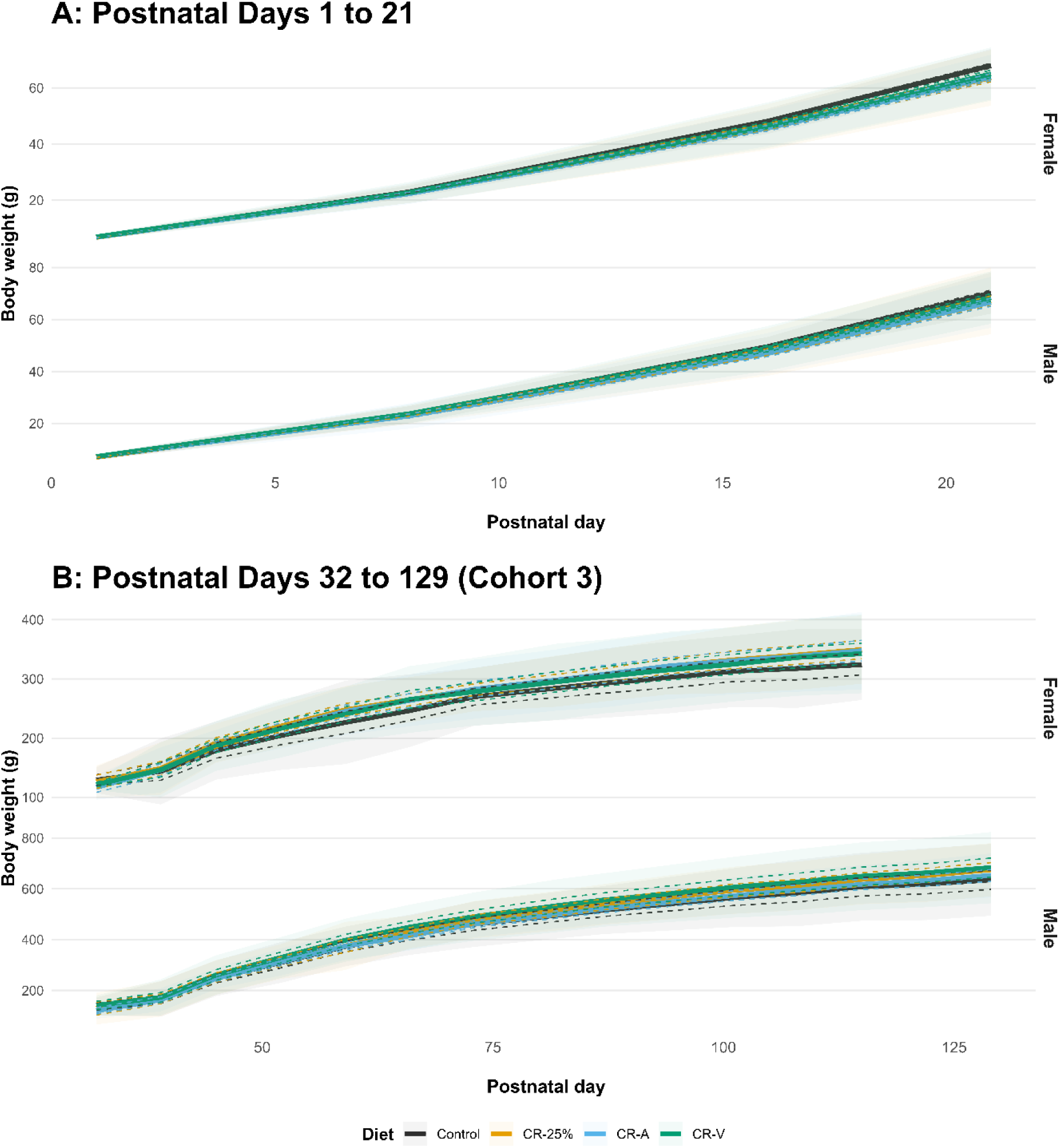
Offspring body weight trajectories across postnatal development in Cohort 3, stratified by Diet and Sex. Body weight (g) is shown for females (top panel) and males (bottom panel) from postnatal (PND) 1-21 (A) and PND 32-129 (B). Solid lines represent mean body weight trajectories for each diet, and shaded regions with dashed boundaries indicate the 95% confidence interval of the mean.

### 2.2 Pup Milestones and Maternal Behaviours

Pup somatic and reproductive milestones were observed daily to assess the impact of the maternal preconception diet. The timing of developmental milestones (fur appearance, pinnae detachment, eye and vaginal opening, and testicular descent was modelled as the day each feature was first observed. ROPE comparisons revealed no meaningful differences between diets across any of the milestones examined.

Additionally, nest quality, maternal care, and pup retrieval performance were assessed to determine whether maternal dietary restriction influenced early caregiving behaviour. Nest construction/quality was rated on a scale from 1 to 4 at 4- and 24-hours following litter standardisation on PND 1. Mean nest rating was good after 4 hours (M = 3.02, SD = 1.14) and 24 hours (M = 3.23, SD = 0.72). No meaningful effect of diet, time point, or their interaction was detected via ROPE comparisons, indicating that maternal preconception diet did not influence nest quality across the first 24 hours postpartum.

Maternal care observations indicated that dams spent most of the time in arch-back/active nursing (M = 39.39%, SD = 12.72%), followed by resting (M = 15.86%, SD = 8.78%). ROPE contrasts revealed that CR-A dams displayed less passive nursing behaviours than control dams [E_m_ = -0.51 [-1.01, -0.10], D_p_ = 98.35, ROPE_p_ = 1.4], while CR-V dams displayed greater out-of-nest activity than control dams [E_m_ = 0.45 [0.16, 0.77], D_p_ = 99.6, ROPE_p_ = 0.8]. No other maternal or non-maternal behaviours differed between the treatment groups, indicating that maternal care was largely unaffected by maternal preconception diet.

In the pup retrieval test, dams had a high probability of retrieving their first pup (M = 0.77, SD = 0.42), whereas the probability of retrieving all pups was lower and below 0.5 (M = 0.44, SD = 0.49). Additionally, the mean latency to retrieve the first pup was 79.8 seconds (SD = 123.42) and 108.02 seconds (SD = 163.02) to retrieve all pups. ROPE contrasts (reported on the logit scale) revealed that CR-A dams were more likely to retrieve all pups than control dams [E_m_ = 1.14 [0.11, 2.36], D_p_ = 96.75, ROPE_p_ = 4.35], whereas no effects were found for the likelihood to retrieve the first pup or for the latencies to retrieve the first or all pups.

### 2.3 Offspring Behaviours

#### 2.3.1 Locomotor Activity

Locomotor activity was assessed using distance travelled and number of zone entries across two testing sessions. Across all offspring, animals travelled an average of 59.5 m (SD = 16.8) and made 275.2 (SD = 89.12) zone entries during the first session, and 44.2 m (SD = 13.46) with 198.2 (SD = 71.29) zone entries during the second session, consistent with habituation across sessions.

ROPE contrasts on the log scale indicated that zone entries were lower in the second session compared to the first for all diet-sex groups [E_m_ = 0.34 [0.26, 0.41], D_p_ = 100, ROPE_p_ = 0], whereas distance travelled [E_m_ = 0.32 [0.27, 0.38], D_p_ = 100, ROPE_p_ = 0], was meaningfully lower in the second session compared for all groups except for female CR-V female offspring, [E_m_ = 0.12 [-0.04, 0.27], D_p_ = 93.6, ROPE_p_ = 8.75]. This indicates that maternal preconception diet selectively reduced locomotion in female offspring.

ROPE contrasts showed no main effect of maternal preconception diet on either distance travelled or zone entries when males and females were considered together. Within sessions, however, sex-specific effects of maternal diet were detected. Pairwise contrasts in the first session, indicated that female offspring from the CR-V group travelled less distance [E_m_ = -0.24 [-0.49, -0.01], D_p_ = 97.9, ROPE_p_ = 2.65], and made fewer zone entries [E_m_ = -0.25 [-0.51, 0.03], D_p_ = 96.15, ROPE_p_ = 3.85] than control female offspring, with effects evident on the log scale.

CR-25% [E_m_ = -0.26 [-0.54, 0.01], D_p_ = 96.5, ROPE_p_ = 3.1] and CR-A [E_m_ = -0.3 [-0.55, - 0.03], D_p_ = 99.15 ROPE_p_ = 1.4] female offspring also made fewer zone entries compared with control female offspring, indicating reduced exploratory activity. No corresponding differences were observed among male offspring. Together, these findings indicate that maternal preconception CR selectively reduces locomotor and exploratory activity in female offspring, without producing global effects across sexes or consistent alterations in habituation.

#### 2.3.2 Running Wheels

Offspring travelled an average of 229.8 m (SD = 145.68) during testing in the running wheel. ROPE contrasts did not reveal any meaningful differences in distance travelled across diets or diet-by-sex pairs, indicating that maternal preconception diet has no impact on offspring voluntary locomotion. Notably, female offspring travelled 319.9 m (SD = 147.30) and male offspring travelled 139.8 m (SD = 68.06). ROPE analysis of distance travelled confirmed that female offspring travelled further than male offspring across all dietary groups (log scale) [E_m_ = 0.83 [0.64, 1.04], D_p_ = 100, ROPE_p_ = 0]. This indicates that whilst preconception diet does not affect voluntary locomotion, sex does cause meaningful differences.

#### 2.3.3 Elevated Plus Maze

Offspring spent an average of 20.7 s (SD = 24.3) in the open arms during testing, entered them 2.2 times (SD = 2.88) and showed a latency of 18.3 s (SD = 32) to first entry. ROPE contrasts did not reveal any meaningful differences between diets on any anxiety-like variables in the EPM in male or female offspring. However, pairwise comparisons revealed that CR-A female offspring travelled less than control female offspring (log scale) [E_m_ = -220.39 [-390.40, 42.39], D_p_ = 99.4, ROPE_p_ = 1.15]. This indicates that maternal preconception diet did not alter the anxiety-like behaviour of offspring, although acute episodes of maternal preconception restriction may reduce overall movement during testing in the EPM in female offspring.

In contrast, robust sex differences were observed. Firstly, ROPE comparisons revealed that female offspring travelled further than male offspring [E_m_ = 314.8 [222.86, 398.47], D_p_ = 99.85, ROPE_p_ = 0.4]. Given this, distance travelled was added as a covariate to the remaining variables of interest. Critically, female offspring entered the open arms more frequently (log scale) [E_m_ = 1.47 [1.02, 1.91], D_p_ = 100, ROPE_p_ = 0], but entered them for the first time later (log scale) [E_m_ = 1.97 [0.66, 3.12], D_p_ = 99.9, ROPE_p_ = 0.35] than male offspring. Additionally, females spent more time in the open arms than males (log scale) [E_m_ = 0.96 [0.38, 1.54], D_p_ = 99.85, ROPE_p_ = 0.4], which was consistent for all diets except CR-A [E_m_ = 0.35 [0.68, 1.32], D_p_ = 74.4, ROPE_p_ = 25.4]. These differences indicate that, regardless of maternal preconception diet, females display lower anxiety-like behaviour during testing in the EPM.

#### 2.3.4 Open Field

Offspring spent an average of 8.1 s (SD = 9.18) in the centre zone, entered it 4.36 (SD = 3.96) times and showed a mean latency of 145.6 s (SD = 146.36) to first entry. ROPE contrasts did not reveal any meaningful differences between diets or across diet-by-sex comparisons for any of these variables. In contrast, sex differences were evident irrespective of diet with ROPE comparisons revealing that female offspring travelled further than males (log scale) [E_m_ = 0.26 [0.18, 0.36], D_p_ = 100, ROPE_p_ = 0]. As such, distance travelled was added as a covariate to the remaining variables of interest. Critically, female offspring spent more time in the middle zone (log scale) [E_m_ = 1.02 [0.6, 1.42], D_p_ = 100, ROPE_p_ = 0], and in the centre zone (log scale) [E_m_ = 2.47 [1.44, 3.45], D_p_ = 100, ROPE_p_ = 0] than male offspring. They also, entered the centre zone more frequently (log scale) [E_m_ = 0.72 [0.38, 1.11], D_p_ = 100, ROPE_p_ = 0] but entered later (log scale) [E_m_ = 2.22 [0.72, 3.66], D_p_ = 99.8, ROPE_p_ = 0.15] than male offspring. Collectively, these differences indicate that, regardless of maternal preconception diet, females display lower anxiety-like behaviour during testing in the open field.

#### 2.3.5 Back Test

Stress reactivity and coping style were assessed in the back test using escape attempts and vocalisations across two sessions. Across all offspring, animals made an average of 12.2 (SD = 2.92) escape attempts and emitted 8.8 (SD = 12.85) vocalisations during testing.

Within-sex group ROPE comparisons indicated that dietary manipulations produced limited, diet-specific effects on escape attempts and vocalisations that depended on both sex and testing session, rather than uniform changes across offspring. Notably, CR-V offspring made more escape attempts than control offspring (log scale) [E_m_ = 0.10 [0.01, 0.20], D_p_ = 97.95, ROPE_p_ = 4.05]. This effect was driven by meaningfully higher escape attempts in CR-V female offspring compared to control females during the first session (log scale) [E_m_ = 0.16 [0.01, 0.33], D_p_ = 96.65, ROPE_p_ = 3.75]. CR-25% female offspring displayed higher escape attempts than control female offspring during the second session (log scale) [E_m_ = 0.17 [0, 0.35], D_p_ = 97.25, ROPE_p_ = 3.55]. No meaningful differences were observed in male offspring in either session.

Additionally, CR-V offspring vocalised less than control offspring overall (log scale) [E_m_ = −0.72 [−1.50, −0.10], D_p_ = 96.05, ROPE_p_ = 3.85]. This reduction was driven by meaningfully lower vocalisations in CR-V female offspring during the second session only (log scale) [E_m_ = −1.25 [−2.47, −0.03], D_p_ = 97.4, ROPE_p_ = 1.5], whereas vocalisations in female offspring during the first session and in male offspring across sessions were not meaningfully different.

CR-A offspring vocalised less than control offspring overall (log scale) [E_m_ = −0.93 [−1.74, −0.11], D_p_ = 98.9, ROPE_p_ = 1.55]. This effect was attributable to meaningfully lower vocalisations in CR-A male offspring compared to control males in both the first session (log scale) [E_m_ = −0.87 [−1.82, 0.13], D_p_ = 96.65, ROPE_p_ = 2.8] and the second session (log scale) [E_m_ = −1.34 [−2.69, −0.11], D_p_ = 98.3, ROPE_p_ = 1.4]. In contrast, female offspring did not differ meaningfully from controls in either session. Despite lower vocalisations, CR-A male offspring made more escape attempts than control males during the second session (log scale) [E_m_ = 0.15 [0.02, 0.31], D_p_ = 96.45, ROPE_p_ = 4.7], with no other CR-A escape contrasts reaching meaningful differences. Together, these findings indicate that maternal preconception dietary restriction produces selective sex-specific alteration in coping, with enhanced active coping responses evident in CR-V and CR-25% female offspring and CR-A male offspring.

#### 2.3.6 Forced Swim Test

Stress-coping strategies were assessed by the amount of time spent floating or swimming during the forced swim test, where offspring typically spent an average of 216.5 seconds (SD = 44.24) floating. ROPE contrasts did not reveal any meaningful differences in floating duration between diets, nor were any diet by sex contrasts meaningfully different. This indicates that coping strategies during forced swimming were not impacted by maternal preconception dietary restriction in either sex.

#### 2.3.7 Sucrose Preference Test

Sucrose preference was compared at 1 and 24 hours after the FST. Mean preference scores were 80.4 % (SD = 16.24) after 1 hour and 88.9 % (SD = 21.61) after 24 hours. ROPE comparisons revealed that maternal preconception dietary restriction did not alter sucrose preference (anhedonic-like behaviour) in offspring following the FST. Additionally, neither sex nor diet-by-sex interactions comparisons revealed any meaningful differences.

#### 2.3.8 Male-Female Interaction Observations

Diet groups were observed for aggressive, male copulatory, dominance, social, non-social, and sociosexual behavioural clusters during individual male-female interactions. Figure 3 shows the counts per session by group for each sexual behavioural cluster observed during male-female interactions, where dietary restriction offspring displayed lower aggressive behaviours and higher male copulatory behaviours than control offspring. In contrast, little variation was observed in the counts per session for social behavioural clusters between groups (see Figure 1 in Supplementary 4 for the time series).

**Figure 3.**
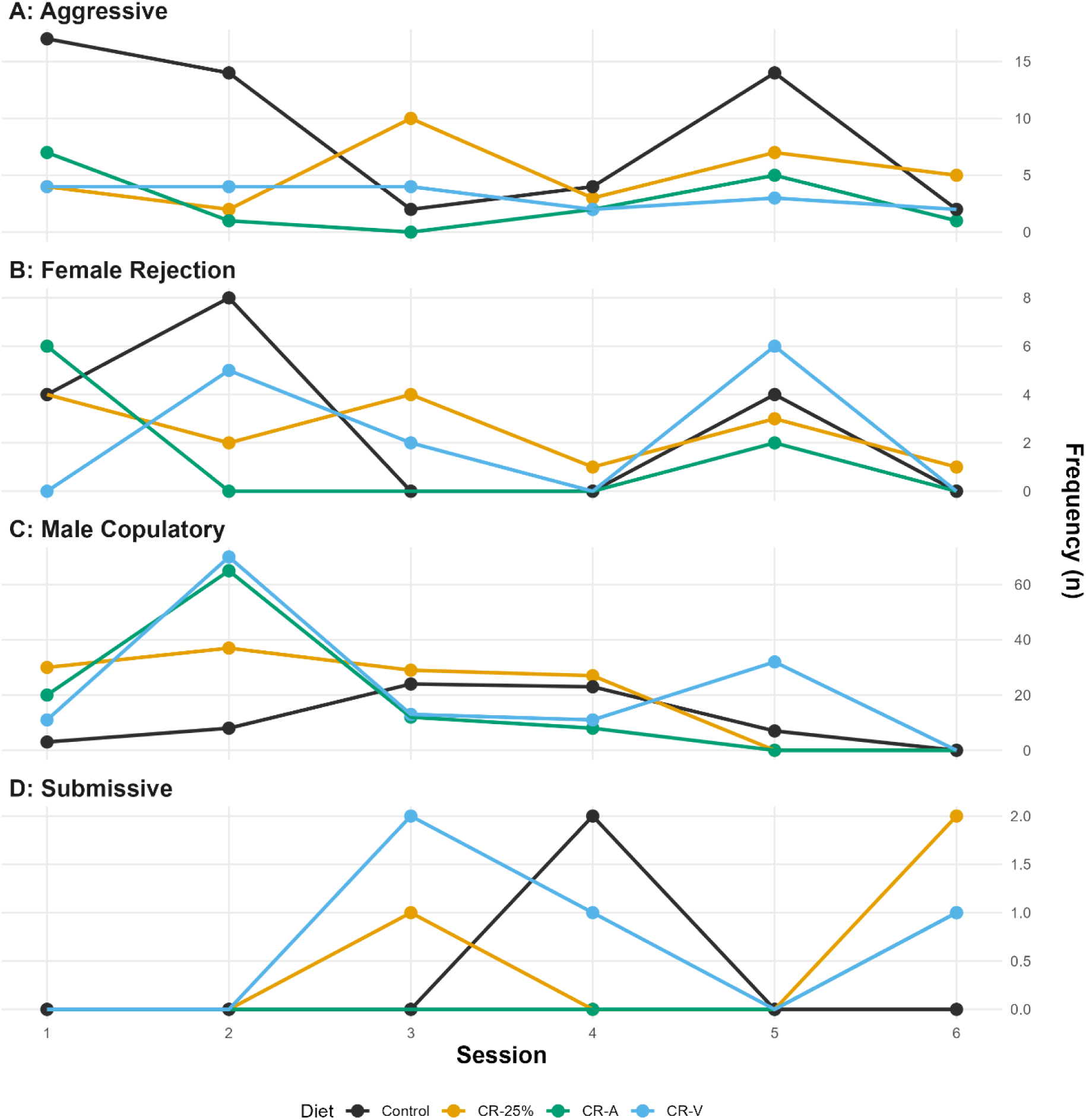
Frequencies of social behaviours between males and females across days by treatment group. Line plots show the observed frequency of behaviours from four functional clusters: Aggressive, Female Rejection, Male Copulatory, and Submissive, recorded over six sessions. Each panel corresponds to one behavioural cluster, with lines representing treatment groups (Control, CR-25%, CR-A, CR-V). Variability in patterns is apparent across both behaviours and groups, particularly in copulatory interactions, where peak frequencies occurred early in the observation period.

As noted, given the fixed time window for observations, the behaviours were analysed using a multinomial model to estimate the probability of observing any one behavioural cluster. Figure 4 shows the estimated probability differences in observing aggressive behaviours per session between diet groups. During the first two sessions of testing, control offspring had a meaningfully higher probability of displaying aggressive behaviours than offspring from any treatment group. In contrast, during the same sessions, the control group showed a meaningfully lower probability of displaying male copulatory behaviours (see Figure 5). All other behavioural clusters exhibited only sporadic differences without consistent effects across sessions and are therefore presented in Supplementary 5-10.

**Figure 4.**
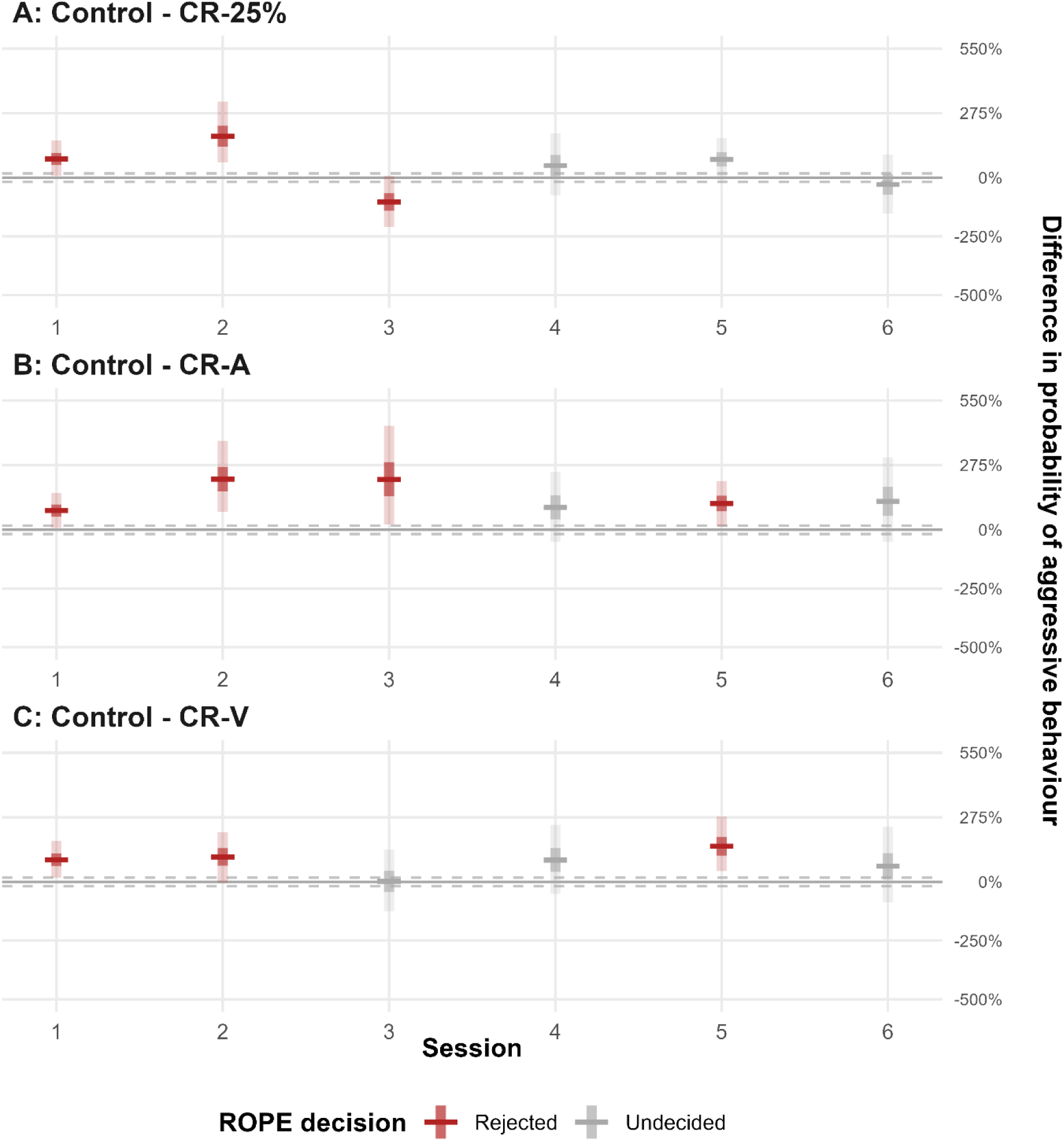
ROPE analysis contrast plots over time for differences in aggressive behaviour. This figure displays the estimated differences in the probability of aggressive behaviour between pairs of treatment groups at six time points (Sessions 1-6). Horizontal lines indicate posterior means, with darker vertical shaded areas representing 50% credible intervals and lighter areas representing 95% credible intervals. ROPE range: 0 ±18%. Positive values indicate a higher probability of aggression in the first group of the contrast compared to the second.

**Figure 5.**
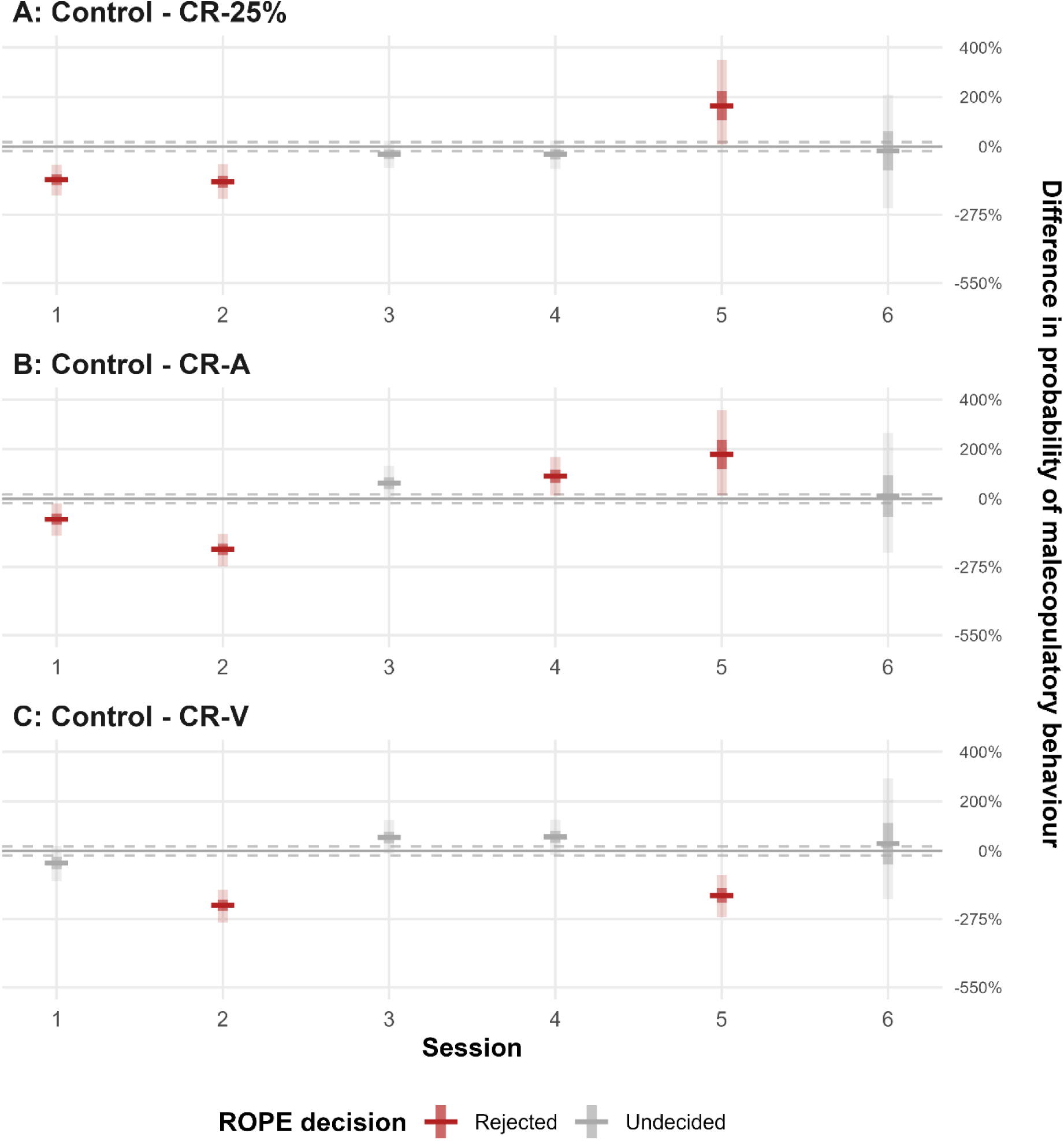
ROPE analysis contrast plots over time for differences in male copulatory behaviour. This figure displays the estimated differences in the probability of male copulatory behaviour between pairs of treatment groups at six time points (Sessions 1-6). Horizontal lines indicate posterior means, with darker vertical shaded areas representing 50% credible intervals and lighter areas representing 95% credible intervals. ROPE range: 0 ±18%. Positive values indicate a higher probability of aggression in the first group of the contrast compared to the second.

##### 2.3.8.1 Male Copulatory / Aggressive Behaviour Bias

Given the pattern of differences between diet and control groups, a follow-up analysis of aggressive and male copulatory behaviours was conducted. To enable a focused comparison, a Bayesian logistic regression model with only two categories was refit: male copulatory and aggressive behaviours. Figure 6 shows the bias between these two behavioural clusters as the posterior distribution of the difference in the probabilities of observing each behaviour.

**Figure 6.**
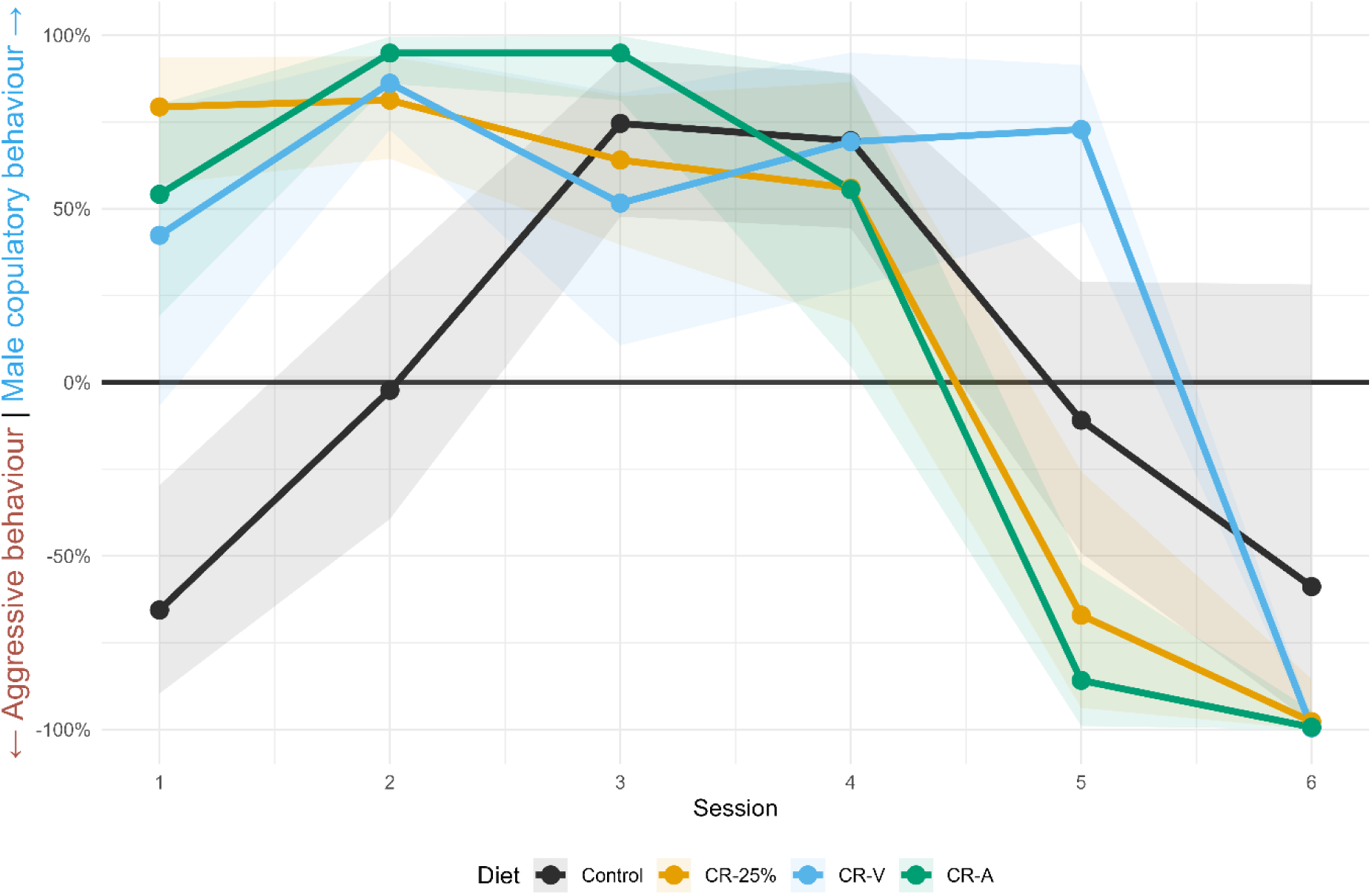
Posterior distributions of the male copulatory bias binary model by group across testing days. The y-axis represents the relative behavioural bias, expressed as a percentage difference between observations of male copulatory and aggressive behaviours. Positive values indicate a relative bias toward male copulatory behaviour, while negative values reflect a bias toward aggressive behaviours. Lines and shaded ribbons show posterior means and 95% credible intervals, respectively, for each experimental group across Sessions 1 to 6.

Notably, the control offspring exhibit a bias toward aggressive behaviours relative to male copulatory behaviours during the first sessions, after which they show no bias between the behaviours in the second session, followed by a shift toward male copulatory behaviours in sessions 3 and 4. Conversely, all offspring from dams exposed to maternal CR displayed a bias toward male copulatory behaviours during the first four sessions, with all restricted offspring except the CR-V offspring shifting to an aggressive behaviour bias for sessions 5 and 6. ROPE contrasts between diets for male copulatory/aggressive behaviour bias (see Figure 7) confirmed that male offspring in all three diet groups showed a meaningful difference in their male copulatory bias in the first 2 sessions of interaction testing compared to control male offspring. This indicates that all maternal preconception diets shifted allocation from aggression to male copulatory behaviours in initial male-female interactions.

**Figure 7.**
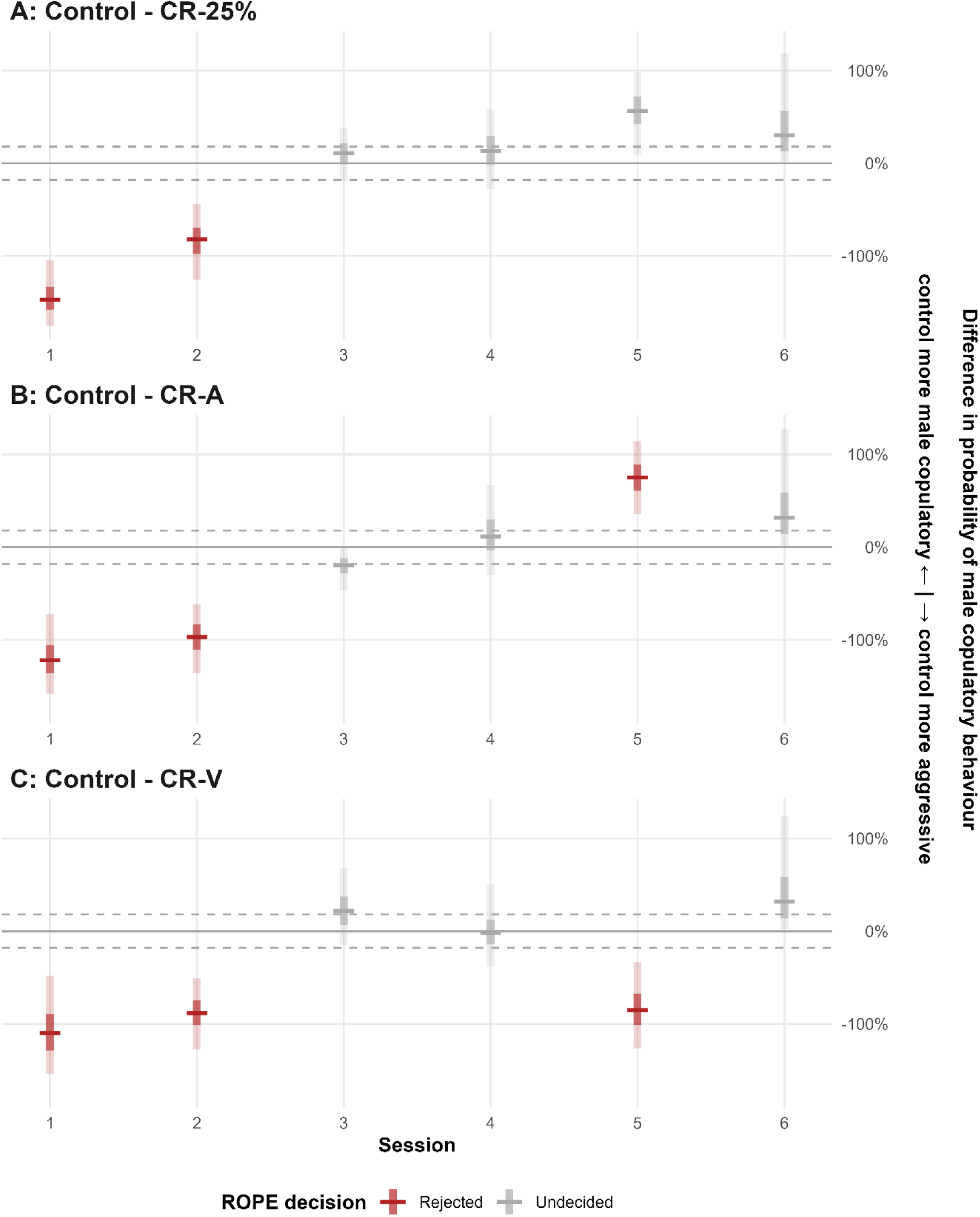
ROPE contrasts estimating group differences in male copulatory bias across testing sessions. This figure displays the estimated differences in male copulatory behaviour bias between pairs of treatment groups across six time points (Session 1 to Session 6). Horizontal lines indicate posterior means, with darker vertical shaded areas representing 50% credible intervals and lighter areas representing 95% credible intervals. ROPE range: 0 ±18%. Positive values indicate a higher probability of male copulatory behaviours relative to aggressive behaviours in the first group of the contrast compared to the second. Conversely, negative values indicate a higher probability of aggressive behaviours relative to male copulatory behaviours in the first group of the contrast compared to the second.

#### 2.3.9 Resident-Intruder Test

When assessing sociality toward an unfamiliar male, male offspring displayed an average frequency of 10.5 (SD = 5.06) submissive, 8.4 (SD = 6.27) mounting, 0.9 (SD = 0.92) dominance, and 4.3 (SD = 9.29) aggressive behaviours. ROPE comparisons (reported on the log scale) between diets revealed that CR-A male offspring exhibited fewer mounts than control male offspring [E_m_ = 1.22 [0.32, 2.16], D_p_ = 99.45, ROPE_p_ = 0.45]. Diet comparisons were not meaningfully different for other behaviours, indicating that submissive, dominant, and aggressive interactions were largely preserved across dietary groups.

##### 2.3.9.1 Faecal Testosterone Metabolites (fTM)

The fTM concentration in faeces prior to and following exposure to the resident-intruder test is shown in Figure 8A below. ROPE contrasts (reported on log scale) revealed that CR-A male offspring displayed a higher concentration of fTM before resident intruder testing compared to control male offspring [E_m_ = 0.44 [0.07, 0.95], D_p_ = 96, ROPE_p_ = 3]. No other diet comparison were meaningfully different within sessions. Notably, fTM concentration increased following exposure to an intruder in the resident-intruder test for control [E_m_ = 0.61 [0.20, 1.00], D_p_ = 100, ROPE_p_ = 1], CR-25% [E_m_ = 1.03 [0.59, 1.46], D_p_ = 100, ROPE_p_ = 0], and CR-A [E_m_ = 0.49 [0, 0.92], D_p_ = 98, ROPE_p_ = 2], but not in CR-V male offspring. This indicates that maternal preconception diet elevated basal fTM prior in CR-A male offspring and suppressed the expected increase following exposure to an intruder in the CR-V offspring.

**Figure 8.**
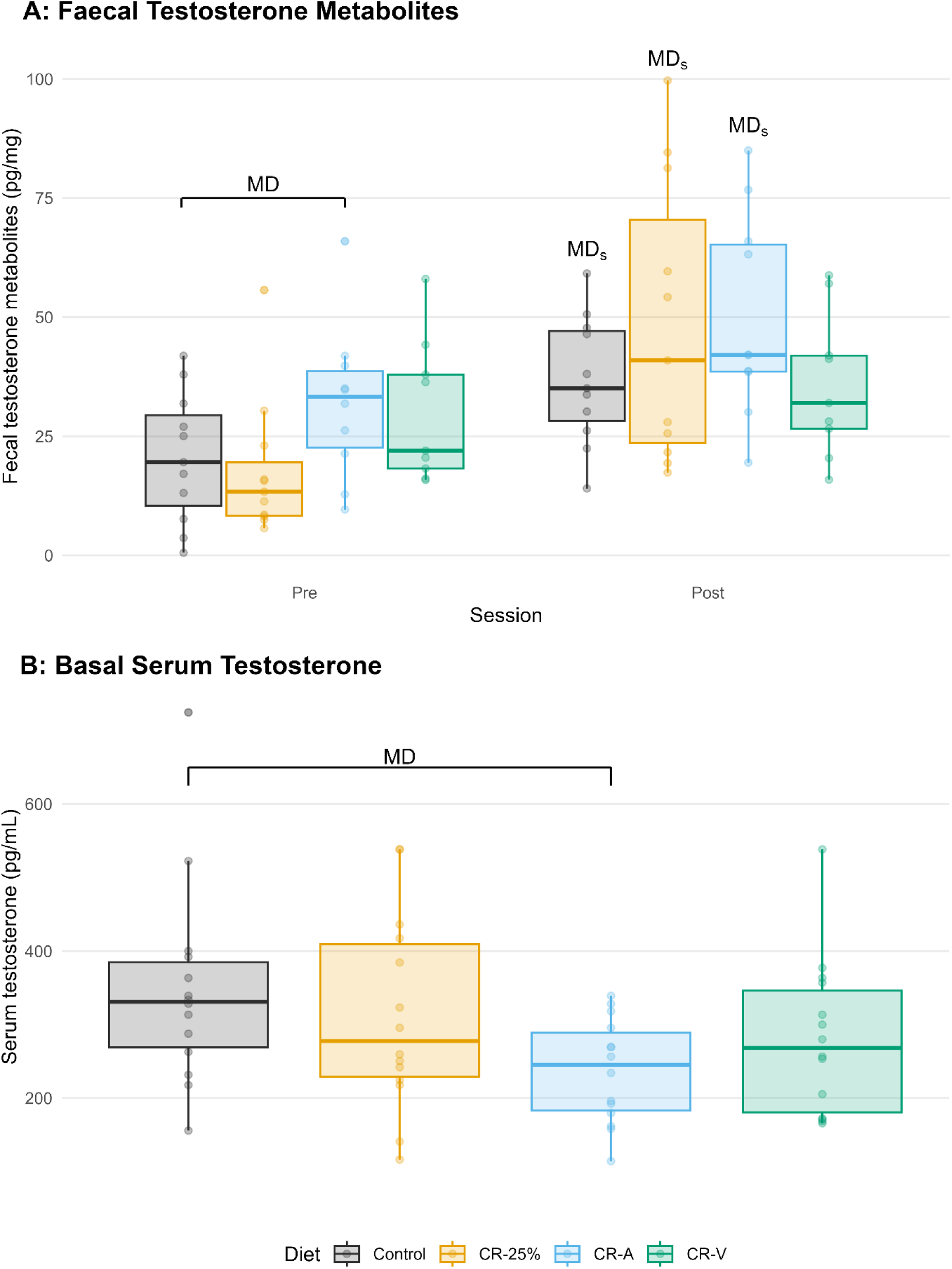
Faecal testosterone metabolite concentration prior to and following resident intruder testing and basal serum testosterone by diet. Boxplots show faecal testosterone metabolites (pg/mg) prior to and following resident intruder testing (A) and basal serum testosterone (pg/mL), (B). Data are presented across the four diet conditions (Control, CR-25%, CR-A, CR-V). Each point represents an individual animal. MD = Meaningfully different. MDS = Meaningful difference relative to the same group’s pre-session measurement.

#### 2.3.10 Basal Serum Testosterone

Basal serum testosterone concentration in males for each diet is shown in Figure 8B. ROPE contrasts (reported on the log scale) indicated that CR-A male offspring had a meaningfully lower testosterone concentration than control male offspring [E_m_ = 0.38 [0.12, 0.64], D_p_ = 99, ROPE_p_ = 1], CR-V male offspring showed a borderline meaningful reduction [E_m_ = 0.21 [-0.06, 0.47], D_p_ = 92.65, ROPE_p_ = 6], and CR-25% male offspring did not differ from controls. This indicates that only acute maternal preconception CR affected basal serum testosterone levels in adult male offspring.

## 3 Discussion

This study comprehensively assessed the effects of multiple maternal CR manipulations administered during preconception on behavioural and neuroendocrine outcomes in male and female adult offspring. The results demonstrate that the preconception developmental window is indeed sensitive to maternal CR, with behavioural and neuroendocrine modifications observed in offspring despite no direct dietary manipulation. Broadly, we observed diet- and sex-specific effects of preconception CR in tests assessing intrinsic exploratory motivation and general activity, and coping behaviour/style. Specifically, we observed a transient reduction in exploratory behaviour in females across all preconception dietary manipulations, particularly in the variable restriction group (CR-V). Diet- and sex-specific effects were also observed, where stable CR (CR-25%) and CR-V adult females and unpredictable food deprivation (CR-A) adult males exhibited a transient active coping style.

Notably, preconception maternal CR, irrespective of the specific regimen, enhanced male offspring sexual behaviour and suppressed aggression toward females. CR-V male offspring failed to show the expected increase in fTM following the resident-intruder test, while basal fTM were increased by preconception CR-A. Moreover, basal serum testosterone was reduced in CR-A offspring, with a similar trend in the CR-V group.

To attribute any behavioural or hormonal changes in offspring to our preconception nutritional manipulations, it was important to exclude any potential effects of differences in dam fecundity, weight, litter size, or pup ratio, as well as pup growth and development. Preconception CR produced clear, though short-term, modifications in dam body weight. Both the CR-25% and CR-V restriction groups lost weight during the 14-day diet phase, though their weights recovered rapidly upon refeeding before mating. Females, across all dietary regimens, conceived and carried litters commensurate with control conspecifics, and offspring postnatal weight, somatic, and reproductive developmental milestones were similarly unaffected. Accordingly, preconception CR did not impair dam fecundity or the early development of progeny.

Maternal behaviour plays a role in the neurodevelopment and behavioural outcomes of offspring, with differences in the quality and quantity of care generating phenotypic differentiation in adulthood [23,26]. Given that nutritional disturbances have been observed to modify maternal caregiving [18,27], it was critical to quantify maternal activities here to contextualise offspring outcomes. Overall, the behavioural profiles of dams do not indicate any consistent or widespread disruption or augmentation of maternal care. All dams, regardless of preconception diet, were equally motivated and capable of constructing adequate nests, suggesting that basic maternal preparation and motivation were intact across diet groups. For the maternal behaviour observations, dams exposed to CR-A displayed less passive nursing, and CR-V dams spent more time out of the nest, consistent with greater environmental monitoring. These modifications were subtle and selective rather than generalised, so that overall maternal care engagement was maintained. Dams in the CR-A group also engaged in more active pup retrieval, suggesting heightened pup responsiveness and goal-directed care giving. Whilst these results reflect subtle shifts in energy management or arousal under preconception dietary restriction, core maternal motivation and caregiving capacity were preserved.

Regarding offspring behavioural alterations resulting from preconception nutritional manipulations, we observed a transient reduction in exploratory motivation in females from all preconception CR groups, as evidenced by fewer zone entries during their first exposure to the locomotion cells. The effect was most substantial in the CR-V females, who also travelled less than controls during the first exposure. This effect did not extend to the second exposure to this test. As the initial exposure to locomotion-based tests can be confounded by anxiety-related inhibition of movement [28], the sessions should be interpreted independently. Accordingly, the hypoactivity of CR female offspring during the initial exposure likely reflects an anxiety-related inhibition of novel exploration, and the group convergence in the second session suggests that baseline locomotion was unaffected by preconception CR once novelty effects subsided (habituation). During testing of state anxiety via the approach-avoidance conflicts elicited by the EPM and OF [29], CR-A females travelled less than control females. This outcome further substantiates the reduction in exploratory-induced locomotion in response to novelty consequent of maternal preconception CR, whilst extending the observation to approach-avoidance tasks. No persistent diet-related differences were observed in voluntary wheel running, demonstrating that intrinsic motivation and spontaneous locomotion were similarly unaffected by our preconception dietary manipulations. Collectively, these findings suggest maternal preconception CR transiently heightens anxiety-driven inhibition of exploration without altering baseline activity.

Regarding coping strategies, diet and sex-specific effects were evident. In the back test, a measure of trait-like coping style [30,31], CR-V female and CR-25% female offspring made more escape attempts in the first session and second, respectively, compared to controls, indicating a transient active coping style [30]. Furthermore, CR-A males made more escape attempts in the second session compared to controls. Conversely, CR-A males vocalized less than controls across both sessions, and CR-V females vocalized less than controls in the second session. Audible squeaks are typically emitted during restraint, handling, or painful procedures and are considered alarm calls signalling immediate discomfort or threat [32]; thus, reduced vocalizations may suggest muted emotional reactivity. Sucrose preference, a measure of anhedonia, was unchanged by preconception dietary manipulations. Overall, maternal CR shaped offspring coping in a sex- and diet-specific manner, with females exposed to stable or unpredictable CR and males exposed to unpredictable food deprivation displaying a more active coping profile.

The most pronounced and consistent effects of maternal preconception CR emerged in the social domain for male offspring. All preconception CR interventions led to a meaningful shift in behavioural strategy during mating. Specifically, preconception dietary manipulations significantly increased copulatory behaviour and reduced aggressive behaviour during the first two male-female pairings compared to the control group. Correspondingly, all CR group male offspring displayed a bias toward male copulatory behaviour rather than aggressive behaviour during the initial pairings, with all groups except the CR-V group shifting to an aggressive bias in later pairings. In contrast, the control group displayed an aggressive bias in the initial and last pairings. Enhanced copulatory efficiency aligns with Govic, Kent, Levay, Hazi, Penman and Paolini [6], who reported improved sexual performance in male offspring of females exposed to severe maternal CR preconception. Our findings extend this work by tracking multiple interactions and incorporating aggression and dominance measures. Collectively, these results indicate that preconception CR promotes a mating-oriented, prosocial phenotype characterised by reduced female-directed aggression.

To profile hormonal mediators of socio-sexual behaviour, we quantified fTM before and after the resident-intruder test and measured basal serum testosterone at euthanasia. Male CR-A offspring showed higher fTM prior to social provocation, consistent with enhanced basal hypothalamic-pituitary-gonadal (HPG) axis activity relative to controls. All groups exhibited increases in fTM following social challenge relative to pre-test levels, except the CR-V offspring, which showed no change, suggesting blunted androgenic responsiveness to social threat. The absence of additional group differences in fTM following intruder exposure corresponds with the behavioural outcomes, as no alterations in aggressive or dominance were observed. In addition to failing to display the expected post-challenge increase in fTM, CR-V male offspring also approached the threshold for lower basal serum testosterone, further supporting attenuated HPG axis activity. Interestingly, CR-A males exhibited elevated basal faecal fTM alongside reduced serum testosterone concentrations measured at a later time point. Faecal testosterone metabolites typically reflect integrated circulating testosterone with an approximate delay of 4 to 6 hours [33]; this temporal offset was considered in our sampling design (faecal: 1500–1700 h; serum: 1100–1400 h) to account for circadian fluctuations of testosterone [33,34]. The divergence between basal faecal and serum testosterone measures in CR-A males may therefore reflect differences in the temporal integration of androgen output versus point estimates of circulating hormone levels, rather than a direct contradiction between measures. Collectively, these findings indicate that distinct patterns of maternal preconception CR differentially program offspring HPG axis regulation, underscoring maternal dietary status as a key determinant of long-term endocrine adaptation.

The suppression of basal serum testosterone observed in CR-V and CR-A offspring is consistent with findings that chronic CR can downregulate the HPG axis, prioritising energy allocation toward somatic maintenance and survival rather than reproduction [35]. Importantly, in the present study, this effect reflects developmental programming following maternal preconception restriction, rather than the consequences of ongoing energy deficit in the offspring. This finding contrasts with previous reports of preconception CR in which elevated testosterone was associated with improved copulatory efficiency [6]. Such discrepancies may reflect methodological differences in both the duration and pattern of restriction (e.g. 3 days in the aforementioned study as opposed to 14 days), as well as differences in the type of restriction employed (fixed versus variable schedules). In addition, the timing of serum testosterone sampling, which was not specified in [6], may contribute to divergent outcomes given the known circadian and context-dependent variability of circulating testosterone. Sexual behaviour is strongly modulated by testosterone [36], and exogenous testosterone reliably reinstates appetitive and consummatory responses in castrated male rats [37]. However, the blunted serum testosterone observed in the CR-A and CR-V offspring, alongside the absence of differences in CR-25% offspring, indicates that peripheral testosterone concentrations alone cannot explain the observed increased copulatory behaviour. Instead, these findings suggest that central neural mechanisms, potentially involving altered androgen sensitivity or organisation of sexually relevant circuits, play a dominant role in mediating the observed behavioural phenotype.

Although peripheral testosterone provides a substrate for androgen receptor activation and local aromatisation, the proximate drivers of male sexual behaviour are centrally mediated, particularly within hypothalamic and mesolimbic circuits such as the medial preoptic area and nucleus accumbens, where dopaminergic signalling plays a key facilitatory role [38]. Conversely, sexual inhibition is mediated by opioid, endocannabinoid, and serotonergic feedback, which act at multiple levels to constrain this excitatory dopaminergic pathway [38]. Within this framework, epigenetic programming of sexually relevant neural circuits by maternal preconception nutritional disturbance represents a plausible mechanism underlying the enhanced sexual behaviour observed in CR-exposed offspring. While direct neurobiological measures were not made in the present study, reduced availability of serotonergic precursors (e.g. tryptophan) during maternal CR could hypothetically program offspring for diminished hypothalamic serotonergic transmission, given the established inhibitory role of serotonin over dopaminergic release. Consistent with this, antagonism of the 5-HT₂C receptor has been shown to increase prefrontal dopamine, a pathway implicated in enhanced sexual motivation [39]. Such neurochemical adaptations could account for heightened sexual engagement in offspring derived from CR-exposed females, without altering physical reproductive milestones. Whether these effects arise from epigenetic changes originating in germline-mediated epigenetic inheritance or from altered placental or hormonal signalling during early development remains unresolved.

This study demonstrates that the preconception period is a sensitive window during which maternal CR can induce nuanced behavioural and neuroendocrine changes in offspring. While maternal care and pup development remained largely unaffected, offspring behavioural outcomes varied by sex, CR pattern, and test context. Female offspring from CR dams exhibited reduced voluntary locomotion and exploratory behaviour in tests of spontaneous locomotion and exploratory behaviour in aversive environments, with some evidence of more adaptive coping strategies in male and female offspring. Male offspring from all CR manipulations exhibited increased copulatory and reduced aggressive behaviours in response to female interaction. Hormonal analyses revealed elevated basal fTM in the CR-A group, yet reduced serum testosterone in both CR-A and CR-V groups. These findings suggest that even in the absence of direct maternal or developmental disruptions, preconception dietary disturbances can subtly program offspring neuroendocrine and behavioural phenotypes. Overall, the pattern of CR was not a major factor. Future work should explore mechanistic pathways and long-term consequences of these early-life nutritional exposures.

## 4 Materials and Methods

### 4.1 Animals

Forty-eight specific pathogen-free, nulliparous female Wistar rats (9 weeks old; 224.40 ± 1.83 g) and 24 male Wistar rats (9 weeks old; 323.21 ± 4.15 g) were procured from Animal Resources Centre (Western Australia, Australia) and allowed to acclimate for 1 week prior to experimentation. During this period, standard rodent chow (Barastoc, Ridley Corporation, Victoria, Australia) and water were available ad libitum. At 12-13 weeks of age, female animals were mated with one of the 24 male Wistar rats, and 320 offspring from 46 dams were used for subsequent experimentation. To support social and sexual behaviour testing, an additional 56 unrelated adult Wistar males and females were used as stimulus animals and housed in same-sex groups. Sample sizes were determined a priori based on prior literature and power calculations. Dams were the experimental unit for preconception dietary manipulation (n = 12 per group). Offspring numbers (12–14 per group per sex) were powered to detect previously reported effects of CR. Additional animals were required for breeding and social interaction assays but were not included in experimental analyses. These animals were required to be experimentally naïve and were used once per relevant assay to avoid prior social experience effects. Adult male breeders and experimental females were group or pair-housed until mating in individually ventilated cages (462 x 403 x 404 mm; l x w x h; Tecniplast Australia, Australia), while offspring and stimulus animals were housed in standard open-top plastic cages (380 x 270 x 150 mm; l x w x h). All animals were provided wood shavings and shredded paper bedding and maintained under controlled temperature (22 +/- 2°C) and lighting conditions (standard 12:12 hr light: dark cycle; lights on at 0700 h for breeding rats, reversed cycle; lights off at 1000 h for offspring and stimulus animals). All procedures were carried out in accordance with the National Health and Medical Research Council of Australia Code of Practice for the Care of Experimental Animals and were approved by La Trobe University Ethics Committee (approval number AEC21038). Additionally, all procedures are reported in accordance with ARRIVE guidelines.

#### Dietary Regimens

Following acclimation, female rats were randomly assigned to four treatment groups (n = 12/group). Dams were randomly allocated to preconception dietary groups with stratification by body weight to ensure comparable baseline weights across groups. Group allocation was known to personnel involved in animal husbandry and animal scheduling. (1) Control: Control females were allowed ad libitum access to food throughout the experimental period; (2) Calorie restriction 25% (CR-25%): the CR-25% group received a constant 25% reduction in daily food intake for 14 days before mating (27.2 ± 1 g/box); (3) Calorie restriction Acute (CR-A): The CR-A group underwent repeated and unpredictable episodes of food deprivation episodes (6, 10, 15, 24 or 48 hr) 14 days before mating; (4) Calorie Restriction Variable (CR-V): the CR-V group received variable, unpredictable level of food restriction, ranging from 25%-75% reduction of food daily for 14 days prior to mating (9 – 27.5 ± 1 g/box). Food intake of the CR-25% and CR-V groups was calculated by averaging the daily intake of all groups over the 48 hrs preceding the start of food restrictions (average daily amount over 48 hrs: 36.3 ± 1 g/box). For the CR-A group, food intake during refeeding periods was measured over the 14-day manipulation and averaged, corresponding to a net 26% restriction. For the CR-25% and CR-V groups, food was delivered daily within ±2 hours of lights out, while food for the CR-A groups was removed or returned at variable times. All restriction groups were reverted to ad libitum access to food following the 14-day regimens. Restriction regimens are shown in Figure 9.

**Figure 9.**
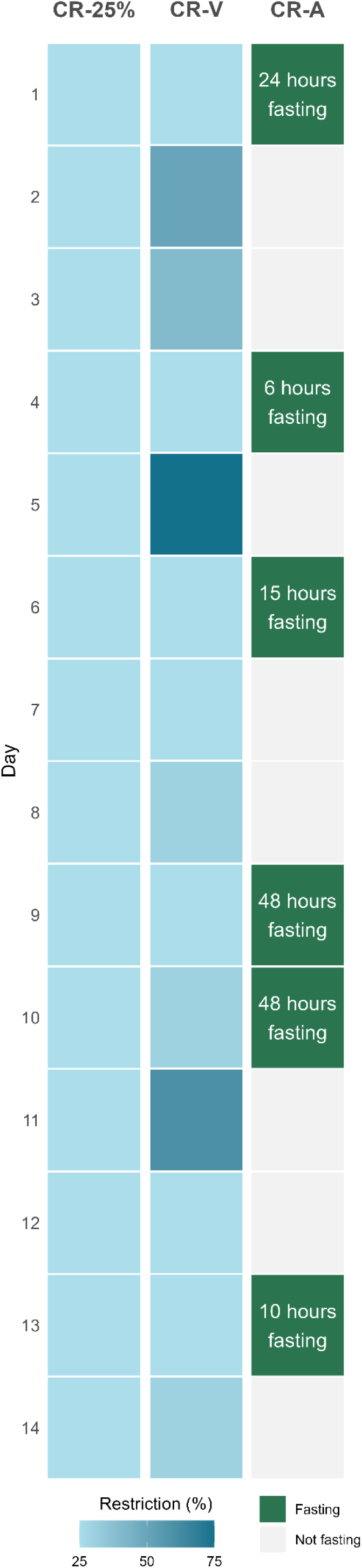
Maternal preconception calorie restriction regimes. Green tiles indicate periods during which CR-A females had no access to food. Blue tiles indicate calorie restricted feeding, with shading reflecting the degree of daily caloric restriction in CR-25% and CR-V females.

### 4.2 Breeding

Twenty-four hrs after CR (CR-25% and CR-V) and 48 hrs after the last food deprivation for the CR-Acute group, male and female breeding rats were paired for 7 days, after which females were individually housed until parturition. To control for paternity, each of the 24 breeding males impregnated two females from two different dietary regimens. The day of parturition was designated as postnatal day (PND) 0.

On PND 1, litters were adjusted to eight pups per dam (4 females, 4 males), except in five cases (one control, two CR-25%, and two CR-V dams) where asymmetrical sex ratios prevented this, and cross-fostering was not viable. If fewer than 8 pups were born, cross-fostering was attempted using excess pups from dams in the same treatment group and born on the same day. This occurred for 1 CR-25% dam. One control dam produced only 7 pups, and two CR-V dams produced 6 pups each, but no cross-fostering was possible. One CR-A female failed to conceive, and another gave birth to a single pup, which was subsequently cannibalised. There were no meaningful differences in litter size or pup sex ratio between dietary groups.

Pup sex was determined on PND1 via anogenital distance and confirmed on PNDs 8, 16, and 21. To minimise interference, dams and pups were weighed on these days only. On PND 21-23, 40 males and 40 females per group (320 offspring) were weaned and pair-housed by treatment group for the remainder of the experiment. Offspring were assigned to three behavioural testing cohorts to avoid carry-over effects from repeated testing. Offspring allocation was constrained by maternal treatment and therefore not randomised across diet groups. Within each maternal treatment group, offspring were assigned to experimental cohorts sequentially based on availability prior to individual identification, with the aim of ensuring representation from each litter within each cohort. The remaining offspring were allocated as needed to balance cohort sizes (Cohort 1 n = 48/sex, 12/group; Cohort 2 n = 56/sex, 12/group; Cohort 3 n = 56/sex, 12/group; Figure 10). All 46 litters were equally represented across the three testing cohorts.

**Figure 10.**
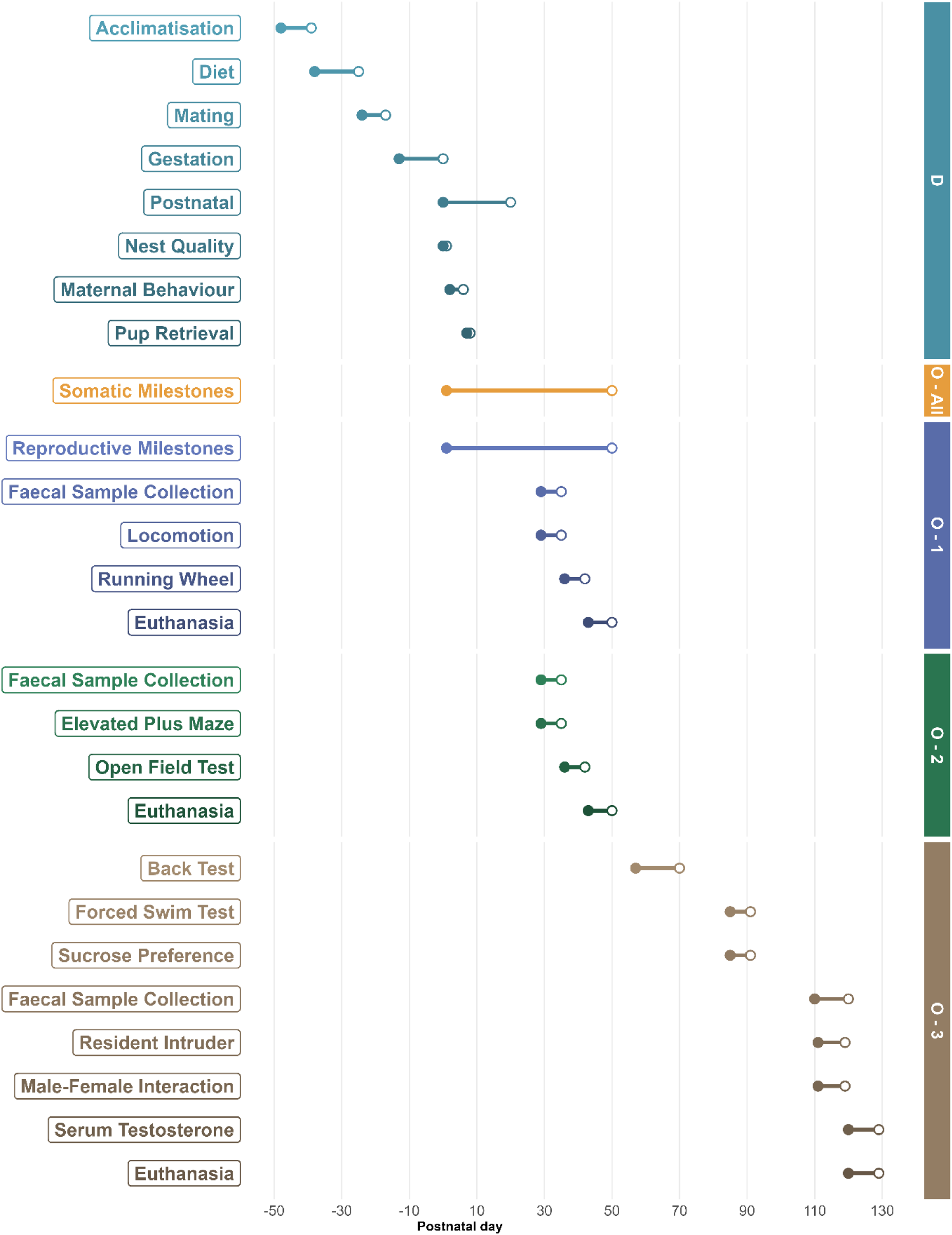
Experimental timeline across postnatal days. Interventions for dams (D) and offspring cohorts (O-All, 1, 2, 3) are shown, along with somatic milestones measured across all three cohorts (O – All). Filled circles = start of testing/sampling, empty circles = end of testing/sampling.

### 4.3 Experiment 1 – Maternal Behaviour and Pup Developmental Milestones

#### 4.3.1 Assessment of Nest Construction/Quality

Nest quality was assessed on PND 1, following litter standardisation. Dams and pups were briefly removed from the cage, along with some of the existing nesting material. Pups were then returned to the cage, and 10 g of shredded paper was scattered across the cage floor. Dams were returned immediately after, with nest quality and maternal contact being scored 4 and 24 hr later. Nest quality was rated on a 5-point scale [40] : 0 (no nest), 1 (poor nest with very little paper used and a flat structure), 2 (fair nest with all paper used but a flat structure), 3 (good nest with all paper used and low walls [<5cm]), and 4 (excellent nest with all paper used and high, compact walls [>5cm]). Maternal contact was scored as dams being on the nest and in contact with the whole litter; on the nest and in contact with part of the litter; or off the nest/off the litter.

#### 4.3.2 Maternal Behaviour Observations

Maternal behaviour was observed from PND 2 to 6 in the home cage under undisturbed conditions. Observations were recorded using Pocket Observer (Noldus, SDR Clinical Tech, Chatswood, NSW, Australia). Cages were arranged prior to parturition to allow full visual inspection of all sides (except behind). Each dam was observed during four 45-min sessions per day: three during the light phase (1000, 1300, 1700 h) and one during the dark phase (0600 h). During each session, behaviours were scored every 3 min (15 observations/session; 4 sessions per day = 60 observations/dam/day), resulting in 300 observations per dam across the 5 days. The following maternal behaviours were coded: arch-back/active nursing, blanket/passive nursing, licking/grooming pups’ heads and anogenital region, retrieval, carrying, mouthing, and stepping. Furthermore, the following non-maternal behaviours were also recorded: self-grooming, eating, drinking, resting, and general out-of-nest behaviour (e.g. rearing, cage exploration, digging). Behavioural definitions are described in more detail in Capone, et al. [41].

#### 4.3.3 Pup Retrieval Test

Pup retrieval was assessed on PND 8 during the light phase (between 1400 and 1600 h). Observations were recorded using Pocket Observer. In this test, dams were temporarily removed from the home cage, and pups were collected and held in a plastic cup for 2 min. Pups were then scattered over the floor on the opposite side of the cage from the nest. The dam was then returned to the cage, and whether the dam received the first pup, the latency to retrieve the first pup, whether the dam retrieved all pups, and the latency to retrieve all pups to the nest were recorded. The test concluded when all pups were returned, or 15 min had elapsed.

#### 4.3.4 Pup Developmental Milestones

Pups were monitored daily for three somatic developmental milestones, consisting of fur development (appearance of dorsal and ventral pigment and fur), eye-opening (both eyes fully opened), and pinnae detachment (both pinnae detached from the cranium).

Reproductive developmental milestones (puberty onset) were assessed in male and female rats from Cohort 1. In female offspring, vaginal opening – defined as complete separation of the membranous sheath covering the vaginal orifice – were assessed daily by visual inspection from PND 25. In male offspring, testis descent – identified by the scrotal purse contacting the testis – was assessed from PND16/18. Assessments continued daily until each milestone was reached or until PND 50.

### 4.4 Experiment 2 – Offspring Behavioural and Neuroendocrine Testing

Offspring were divided into three testing cohorts and exposed to designated test batteries (Figure 10). Cohorts were grouped by behavioural domain (i.e. locomotion, anxiety-like behaviour, and depression-like/coping/socio-sexual behaviour). Behavioural testing was conducted using animal identification numbers that did not indicate experimental group, such that the experimenters conducting testing were not aware of group allocation at the time of data collection. Outcome measures were derived from predefined behavioural metrics. Male offspring underwent the full suite of behavioural tests, whereas female offspring did not undergo sexual behaviour observations or the resident-intruder paradigm. All behavioural testing was conducted during the first 6 hours of the dark portion of the light: dark cycle and included a minimum 30 min habituation period to the testing conditions, where applicable. Exceptions to this habituation included the running wheels and back test. Testing was conducted under standard lighting (∼200 lx of light) except for the male-female observations, which occurred under red lights. All mazes/testing chambers were cleaned with 80% ethanol between tests. Where required, tests were recorded using overhead closed-circuit cameras and analysed with Ethovision XT tracking software (Noldus, SDR Clinical Tech, Chatswood, NSW, Australia) for tasks such as the elevated plus maze and open field. Other tasks, including the back test, forced swim test, and resident-intruder test, were coded offline using Observer XT (Noldus), while male-female interactions were live-coded using Pocket Observer.

Animals were euthanised by gradual-fill carbon dioxide (CO2) inhalation at the conclusion of the designated test battery, in accordance with approved animal ethics protocols. CO2 was used exclusively for euthanasia and not as an anaesthetic agent. No anaesthetic agents were administered at any stage of the study.

#### 4.4.1 Cohort 1 Offspring

##### 4.4.1.1 Locomotor Activity

Rats were placed in automated photocell activity chambers (43 x 43 x 30.5 cm; Med Associates Inc, VT, USA), equipped with 16-beam infrared arrays located on the X, Y, and Z axes. Spontaneous locomotor activity was recorded in two 1 hr sessions conducted 24 hrs apart. The chamber was virtually divided into four equal zones (a two-by-two grid). All variables, except vertical count and vertical time, generated by the Activity Monitor Software (Med Associates Inc, VT, USA) were used for analysis. Both vertical measures were removed due to inconsistency in measuring vertical movement between sexes.

##### 4.4.1.2 Running Wheels

Rats were individually placed in cages containing automated running wheels (35.56 cm in diameter; Lafayette Instrument, Indianapolis, USA) for a 2-hr voluntary running period. The stainless-steel wheels featured 1.59 mm rods spaced 7.94 mm apart. Total distance (cm) travelled was measured via Scurry Activity Monitoring Software (Lafayette Instrument, Indianapolis, USA). Food and water were available throughout testing, and animals were returned to their home cage afterwards.

#### 4.4.2 Cohort 2 Offspring

##### 4.4.2.1 Elevated Plus Maze

In the elevated plus maze (EPM), rats were placed in the centre of a cross-shaped maze consisting of two open and two closed arms (50 x 12 cm; l x w; 40 cm arm walls), elevated 50 cm from the ground. Rats were placed in the centre of the maze facing an open arm and allowed to freely explore the maze for 5 min. Duration and frequency of entries into the open and closed arms of the maze, arm transition frequency, latency to enter the arms, and total distance travelled were recorded and served as indices of anxiety-like behaviour. Zone entry was operationally defined as having two paws in the zone.

##### 4.4.2.2 Open Field

The open field (OF) test was conducted in a grey opaque arena (120 x 120 x 50 cm; l x w x h). Rats were placed facing a wall and allowed to explore for 10 min. Offline analysis divided the arena into concentric central (48 x 48cm), middle (90 x 90cm), and outer zones. Entry into the middle and centre zones was operationally defined as two paws. The total distance travelled, the latency, frequency, and time spent in the middle and centre zones, and the zone transfer frequency were calculated. Longer latencies to enter the centre zone, fewer entries into, and less time spent in the centre zone of the OF served as indices of anxiety-like behaviour.

#### 4.4.3 Cohort 3 Offspring

##### 4.4.3.1 Back Test

Rats were gently restrained in a supine position for 1 min by grasping behind the shoulders and tail base. Rats were exposed to two sessions separated by 7 days. Escape attempts, defined as “wiggling” or movement of the limbs, and vocalisations were recorded by two observers blind to the treatment group. Average scores for the two raters were calculated for both variables. Interrater reliability was substantial (κ = 0.76, p < 0.01), indicating consistent ratings between observers. For the analysis, escape attempts with and without biting were collapsed.

##### 4.4.3.2 Forced Swim Test

The forced swim test consisted of a 10-minute habituation session and a 5-minute test session 24-hrs later in a water-filled Perspex cylinder (20 cm wide × 50 cm high, 30 cm water depth, 23 ± 2°C). During the test session, the latency to immobility/floating, swimming/climbing, and diving, and the frequency and time spent engaged in these activities, were recorded. Reduced immobility, or increased latency to immobility indicated reduced depression-like behaviour and/or a more active coping strategy [42,43]. Coding was performed offline by two blinded observers, and the average of the two raters was calculated for each variable. Interrater reliability was high (κ = 0.86, p < 0.01), indicating consistent ratings between observers.

##### 4.4.3.3 Sucrose Preference Test

The sucrose preference test assessed anhedonia using a two-bottle free-choice paradigm conducted over 48 hours following the forced swim test. Pair-housed animals were given access to two drinking bottles in their home cage: one with water and one with a 2% sucrose solution. Bottles were switched following 24 hr to control for side bias. Sucrose preference was calculated as a percentage of sucrose intake over total fluid intake 1 and 24 hr following the test session of the FST.

##### 4.4.3.4 Male-Female Interaction Observations

For the male-female interaction observations, males were pair-housed with an unfamiliar female conspecific. Females were 35-42% of the weight of males. Estrous cyclicity was not tracked; however, behaviour was sampled across the full estrous cycle (4–5 days in Wistar rats; [44].

Observations occurred under undisturbed red lighting conditions in the colony room across six 45-min observation sessions across five consecutive days. On the first day, pairing occurred 1-2 hrs following lights out, and observations commenced approximately 10 mins post-pairing (pairing session).

Subsequent sessions occurred between 13:30 and 16:00 hrs. Behaviour was sampled every 90 sec for 10 sec (30 observations/session). In this way, each male-female interaction pair was observed 180 times across the 5 days. The presence of the following behaviours were coded [45–47]: male copulatory behaviours (mounts, intromissions, ejaculations); female rejection behaviour (nose-off, posturing); aggressive behaviour (clinch attack, attack, pushing/boxing, lateral threat, kicking, upright standing); dominance behaviour (pinning, walkover); submissive behaviour (fleeing); social behaviour (sniffing, allogrooming); sociosexual behaviour (anogential sniffing, following/chasing); non-social behaviour (self-grooming, resting, cage-exploration, rearing, eating/drinking). The clusters for these behaviours can be found in Supplementary 1.

##### 4.4.3.5 Resident-Intruder Test

The male-female interaction pairings also served to establish male home cage territoriality for the resident-intruder test. After 7-10 days of pair housing, females were removed at least 1 hr before a novel male intruder was introduced to the home cage for a total duration of 15 min. Experimental rats were ∼36 - 37% heavier than intruders. Ethologically related behaviours were aggregated to create composite behavioural scores [45,48]. The aggregates and their associated behaviours included: aggressive behaviours: biting, clinch attack, lateral attack, fighting; dominance behaviours: walk over, walk under, mount; submissive behaviours: move away, fleeing. Observers were blinded to treatment group, and the average of the two raters was calculated for each variable. Inter-rater reliability between two independent coders was good to excellent (κ = 0.62, p < 0.01). Behavioural scores were calculated as the mean of both coders’ ratings for all subsequent analyses.

##### 4.4.3.6 Faecal Testosterone Metabolites (fTM)

Faecal samples were collected four hours after resident-intruder testing to capture peak excretion of testosterone metabolites, reflecting circulating hormone changes with a delay of approximately 4-6 hours during the active (dark) phase in rodents [33]. Baseline samples were collected at the same time of day, approximately 10 days prior to testing and before pair housing with females, to control for circadian and housing-related influences on hormone levels. Faecal pellets were collected directly from the anus into sterile tubes immediately upon handling.

Testosterone was extracted using a steroid solid extraction protocol (Invitrogen, Thermo Fisher Scientific, Australia), and Testosterone was determined by a solid-phase monoclonal, antibody-based, competitive Enzyme-Linked Immunosorbent Assay (EIATESX10; Invitrogen, Thermo Fisher Scientific, Australia) in accordance with the manufacturer’s instructions. Samples were thawed, incubated with the dissociation reagent, and diluted with Assay buffer to a minimum of 1:36 prior to plating in duplicate in antibody-coated microplate wells and incubated for 120 min at room temperature with testosterone conjugate and testosterone antibody. Following the incubation period, the microplate was decanted and washed four times; 100 μL of tetramethylbenzidine solution was added to each well and incubated at room temperature for 30 min. After incubation, colour development was stopped, and the optical density of the wells was measured at 450 nm with an EnSpireTM Multilabel Reader 2300 (PerkinElmer, Wallac Oy, Mustionkatu 6, FI-20750, Turku, Finland). Testosterone levels were calculated using GainData ELISA data calculator (Arigo Biolaboratories). Intra-assay and inter-assay coefficients of variation were 5.33% and 8.57%, respectively. The detection limit of the assay was 7.81 pg/mL.

##### 4.4.3.7 Serum Testosterone ELISA

Serum samples were collected at the time of euthanasia for Cohort 3. Testosterone was determined by a solid-phase monoclonal, antibody-based, competitive Enzyme-Linked Immunosorbent Assay (EIATESX10; Invitrogen, Thermo Fisher Scientific, Australia) in accordance with the manufacturer’s instructions (same procedure as above).

### 4.5 Data and Statistical Analysis

Data analysis was conducted with group identities visible, as group labels were required for statistical analysis. Animals were excluded from specific analyses only where predefined technical or welfare-related criteria were met (e.g. Equipment malfunction, incomplete task engagement, or data acquisition failure). No animals were excluded based on treatment group or outcome. Final sample sizes, including details of exclusions for each test, are specified by test in Supplementary 2.

Statistical analysis of behavioural data was conducted using Bayesian generalised linear regression models, including mixed-effects regression models using subject as the random effect and default priors unless otherwise specified. Model diagnostic criteria included Rhat, Effective Sample Size (ESS) and Leave-one-out Cross Validation [49]. Convergence was accepted when Rhat values were within (0.99, 1.01), ESS > 400, and all k values from cross-validation were < 0.7 [50–52]. Model fit was also visually assessed via posterior predictive checks to ensure that the model could adequately reproduce the observed data patterns. Unless otherwise specified, all models converged with Rhat, ESS and k values within acceptable ranges. Model specifications including family and predictor as well as information regarding non-standard ROPE decisions can be found in Supplementary 2. All prior specifications, regression summaries, diagnostic results, posterior predictive checks, and ROPE bounds can be found in the online repository linked in the Supplementary 3.

Experimental effects were assessed using the Sequential Effect Existence and Significance Testing (SEXIT) framework using the Region of Practical Equivalence (ROPE). For aggregated values, median effect size (EM), 95% highest density interval (HDI), probability of direction (Dp), and proportion of the posterior within a region of practical equivalence (ROPEp) were reported [53]. Meaningful differences were inferred when the probability of the direction of the effect was greater than 95% and less than 5% of the posterior of the effect was contained within the ROPE. Equivalence was accepted where 100% of the distribution was contained within the ROPE; otherwise, the equivalence of the effect was considered undecided. Critically, ROPE range measurements follow recommendations by Kruschke [54], except for weight comparisons, which use 10% of the mean body weight for the sample of interest, as this is a clinically relevant effect size for weight change [55]. Four requisite statistics to indicate the existence of an effect were reported: Median estimate and 95% highest density interval (HDI) of the effect: EM (HDI lower bound, HDI upper bound), the probability of direction of the effect (Dp), and proportion of the effect inside a region of practical equivalence (ROPEp), for example [Em= (), D_p_ = ROPE_p_ = ]. Only meaningful differences will be reported in-text, whereas full ROPE effect summary tables can be found in the online summaries linked in the Supplementary 3.

## Author Contributions

JP, AG, AH, and HN conceptualised and designed the study. AH and AG were responsible for acquiring funding and obtaining the resources required for the project. AG, AH, HN, and MC, designed the methodology and collected the data. MDZ, AG, AH, EAL, HN, TGJ, and MC analysed and interpreted the data, while MDZ visualised the data. MDZ, AH, EAL, and AG wrote the original draft, while AG, EAL, MDZ, AH, HN, MC, TGJ, and JP reviewed and edited the manuscript. All authors have read and agreed to the final version of the manuscript.

## Data Availability Statement

All data supporting the findings of this study are available within the paper and its Supplementary Information. Data is available at https://www.epigenes.com.au/our-research/maternal-preconception-dietary-restriction

## References

1. Pena-Villalobos, I.; Otarola, F.A.; Arancibia, D.; Sabat, P.; Palma, V. Prenatal caloric restriction adjusts the energy homeostasis and behavior in response to acute and chronic variations in food availability in adulthood. J Comp Physiol B 2023, 193, 677–688, doi:10.1007/s00360-023-01520-6.

2. Pico, C.; Palou, M.; Priego, T.; Sanchez, J.; Palou, A. Metabolic programming of obesity by energy restriction during the perinatal period: different outcomes depending on gender and period, type and severity of restriction. Front Physiol 2012, 3, 436, doi:10.3389/fphys.2012.00436.

3. Zhang, Q.; Xiao, X.; Zheng, J.; Li, M.; Yu, M.; Ping, F.; Wang, T.; Wang, X. A Maternal High-Fat Diet Induces DNA Methylation Changes That Contribute to Glucose Intolerance in Offspring. Front Endocrinol (Lausanne) 2019, 10, 871, doi:10.3389/fendo.2019.00871.

4. Akitake, Y.; Katsuragi, S.; Hosokawa, M.; Mishima, K.; Ikeda, T.; Miyazato, M.; Hosoda, H. Moderate maternal food restriction in mice impairs physical growth, behavior, and neurodevelopment of offspring. Nutr Res 2015, 35, 76–87, doi:10.1016/j.nutres.2014.10.014.

5. Govic, A.; Bell, V.; Samuel, A.; Penman, J.; Paolini, A.G. Calorie restriction and corticosterone elevation during lactation can each modulate adult male fear and anxiety-like behaviour. Horm Behav 2014, 66, 591–601, doi:10.1016/j.yhbeh.2014.08.013.

6. Govic, A.; Kent, S.; Levay, E.A.; Hazi, A.; Penman, J.; Paolini, A.G. Testosterone, social and sexual behavior of perinatally and lifelong calorie restricted offspring. Physiol Behav 2008, 94, 516–522, doi:10.1016/j.physbeh.2008.03.007.

7. Monteiro, S.; Nejad, Y.S.; Aucoin, M. Perinatal diet and offspring anxiety: A scoping review. Transl Neurosci 2022, 13, 275–290, doi:10.1515/tnsci-2022-0242.

8. Boersma, G.J.; Bale, T.L.; Casanello, P.; Lara, H.E.; Lucion, A.B.; Suchecki, D.; Tamashiro, K.L. Long-term impact of early life events on physiology and behaviour. J Neuroendocrinol 2014, 26, 587–602, doi:10.1111/jne.12153.

9. Fleming, T.P.; Watkins, A.J.; Velazquez, M.A.; Mathers, J.C.; Prentice, A.M.; Stephenson, J.; Barker, M.; Saffery, R.; Yajnik, C.S.; Eckert, J.J.;, et al. Origins of lifetime health around the time of conception: causes and consequences. Lancet 2018, 391, 1842–1852, doi:10.1016/S0140-6736(18)30312-X.

10. Stephenson, J.; Heslehurst, N.; Hall, J.; Schoenaker, D.; Hutchinson, J.; Cade, J.E.; Poston, L.; Barrett, G.; Crozier, S.R.; Barker, M.;, et al. Before the beginning: nutrition and lifestyle in the preconception period and its importance for future health. Lancet 2018, 391, 1830–1841, doi:10.1016/S0140-6736(18)30311-8.

11. Jahan-Mihan, A.; Leftwich, J.; Berg, K.; Labyak, C.; Nodarse, R.R.; Allen, S.; Griggs, J. The Impact of Parental Preconception Nutrition, Body Weight, and Exercise Habits on Offspring Health Outcomes: A Narrative Review. Nutrients 2024, 16, doi:10.3390/nu16244276.

12. Li, M.; Francis, E.; Hinkle, S.N.; Ajjarapu, A.S.; Zhang, C. Preconception and Prenatal Nutrition and Neurodevelopmental Disorders: A Systematic Review and Meta-Analysis. Nutrients 2019, 11, doi:10.3390/nu11071628.

13. Venter, E.; Zandberg, L.; Venter, P.V.; Smuts, C.M.; Kruger, H.S.; Baumgartner, J. Female rats consuming an iron and omega-3 fatty acid deficient diet preconception require combined iron and omega-3 fatty acid supplementation for the prevention of bone impairments in offspring. J Dev Orig Health Dis 2024, 15, e6, doi:10.1017/S2040174424000102.

14. Watkins, A.J.; Wilkins, A.; Cunningham, C.; Perry, V.H.; Seet, M.J.; Osmond, C.; Eckert, J.J.; Torrens, C.; Cagampang, F.R.; Cleal, J.;, et al. Low protein diet fed exclusively during mouse oocyte maturation leads to behavioural and cardiovascular abnormalities in offspring. J Physiol 2008, 586, 2231–2244, doi:10.1113/jphysiol.2007.149229.

15. Dudele, A.; Lund, S.; Jessen, N.; Wegener, G.; Winther, G.; Elnif, J.; Frische, S.; Wang, T.; Mayntz, D. Maternal protein restriction before pregnancy reduces offspring early body mass and affects glucose metabolism in C57BL/6JBom mice. J Dev Orig Health Dis 2012, 3, 364–374, doi:10.1017/S2040174412000347.

16. Duko, B.; Mengistu, T.S.; Stacey, D.; Moran, L.J.; Tessema, G.; Pereira, G.; Bedaso, A.; Gebremedhin, A.T.; Alati, R.; Ayonrinde, O.T.;, et al. Associations between maternal preconception and pregnancy adiposity and neuropsychiatric and behavioral outcomes in the offspring: A systematic review and meta-analysis. Psychiatry Res 2024, 342, 116149, doi:10.1016/j.psychres.2024.116149.

17. Mortensen, E.L.; Wang, T.; Malte, H.; Raubenheimer, D.; Mayntz, D. Maternal preconceptional nutrition leads to variable fat deposition and gut dimensions of adult offspring mice (C57BL/6JBom). Int J Obes (Lond) 2010, 34, 1618–1624, doi:10.1038/ijo.2010.91.

18. Levay, E.A.; Paolini, A.G.; Govic, A.; Hazi, A.; Penman, J.; Kent, S. Anxiety-like behaviour in adult rats perinatally exposed to maternal calorie restriction. Behav Brain Res 2008, 191, 164–172, doi:10.1016/j.bbr.2008.03.021.

19. Ramirez-Lopez, M.T.; Vazquez, M.; Bindila, L.; Lomazzo, E.; Hofmann, C.; Blanco, R.N.; Alen, F.; Anton, M.; Decara, J.; Arco, R.;, et al. Maternal Caloric Restriction Implemented during the Preconceptional and Pregnancy Period Alters Hypothalamic and Hippocampal Endocannabinoid Levels at Birth and Induces Overweight and Increased Adiposity at Adulthood in Male Rat Offspring. Front Behav Neurosci 2016, 10, 208, doi:10.3389/fnbeh.2016.00208.

20. Fisch, J.; Feistauer, V.; de Moura, A.C.; Silva, A.O.; Bollis, V.; Porawski, M.; Almeida, S.; Guedes, R.P.; Barschak, A.G.; Giovenardi, M. Maternal feeding associated to post-weaning diet affects metabolic and behavioral parameters in female offspring. Physiol Behav 2019, 204, 162–167, doi:10.1016/j.physbeh.2019.02.026.

21. Smith, K.E.; Pollak, S.D. Early life stress and development: potential mechanisms for adverse outcomes. J Neurodev Disord 2020, 12, 34, doi:10.1186/s11689-020-09337-y.

22. Sheng, J.A.; Handa, R.J.; Tobet, S.A. Evaluating different models of maternal stress on stress-responsive systems in prepubertal mice. Front Neurosci 2023, 17, 1292642, doi:10.3389/fnins.2023.1292642.

23. Menard, J.L.; Champagne, D.L.; Meaney, M.J. Variations of maternal care differentially influence ’fear’ reactivity and regional patterns of cFos immunoreactivity in response to the shock-probe burying test. Neuroscience 2004, 129, 297–308, doi:10.1016/j.neuroscience.2004.08.009.

24. Azizi, N.; Roshan-Milani, S.; MahmoodKhani, M.; Saboory, E.; Gholinejad, Z.; Abdollahzadeh, N.; Sayyadi, H.; Chodari, L. Parental pre-conception stress status and risk for anxiety in rat offspring: specific and sex-dependent maternal and paternal effects. Stress 2019, 22, 619–631, doi:10.1080/10253890.2019.1619075.

25. Winther, G.; Eskelund, A.; Bay-Richter, C.; Elfving, B.; Muller, H.K.; Lund, S.; Wegener, G. Grandmaternal high-fat diet primed anxiety-like behaviour in the second-generation female offspring. Behav Brain Res 2019, 359, 47–55, doi:10.1016/j.bbr.2018.10.017.

26. Champagne, F.A.; Curley, J.P. Epigenetic mechanisms mediating the long-term effects of maternal care on development. Neurosci Biobehav Rev 2009, 33, 593–600, doi:10.1016/j.neubiorev.2007.10.009.

27. McGuire, M.K.; Pachon, H.; Butler, W.R.; Rasmussen, K.M. Food restriction, gonadotropins, and behavior in the lactating rat. Physiol Behav 1995, 58, 1243–1249, doi:10.1016/0031-9384(95)02018-7.

28. Miller, C.K.; Halbing, A.A.; Patisaul, H.B.; Meitzen, J. Interactions of the estrous cycle, novelty, and light on female and male rat open field locomotor and anxiety-related behaviors. Physiol Behav 2021, 228, 113203, doi:10.1016/j.physbeh.2020.113203.

29. Calhoon, G.G.; Tye, K.M. Resolving the neural circuits of anxiety. Nat. Neurosci. 2015, 18, 1394–1404, doi:10.1038/nn.4101.

30. Bardi, M.; Rhone, A.P.; Franssen, C.L.; Hampton, J.E.; Shea, E.A.; Hyer, M.M.; Huber, J.; Lambert, K.G. Behavioral training and predisposed coping strategies interact to influence resilience in male Long-Evans rats: implications for depression. Stress 2012, 15, 306–317, doi:10.3109/10253890.2011.623739.

31. Hawley, D.F.; Bardi, M.; Everette, A.M.; Higgins, T.J.; Tu, K.M.; Kinsley, C.H.; Lambert, K.G. Neurobiological constituents of active, passive, and variable coping strategies in rats: integration of regional brain neuropeptide Y levels and cardiovascular responses. Stress 2010, 13, 172–183, doi:10.3109/10253890903144621.

32. Packheiser, J.; Soyman, E.; Paradiso, E.; Michon, F.; Ramaaker, E.; Sahin, N.; Muralidharan, S.; Wohr, M.; Gazzola, V.; Keysers, C. Audible pain squeaks can mediate emotional contagion across pre-exposed rats with a potential effect of auto-conditioning. Commun Biol 2023, 6, 1085, doi:10.1038/s42003-023-05474-x.

33. Auer, K.E.; Kussmaul, M.; Mostl, E.; Hohlbaum, K.; Rulicke, T.; Palme, R. Measurement of Fecal Testosterone Metabolites in Mice: Replacement of Invasive Techniques. Animals (Basel) 2020, 10, doi:10.3390/ani10010165.

34. Chacon, F.; Cano, P.; Jimenez, V.; Cardinali, D.P.; Marcos, A.; Esquifino, A.I. 24-hour changes in circulating prolactin, follicle-stimulating hormone, luteinizing hormone, and testosterone in young male rats subjected to calorie restriction. Chronobiol Int 2004, 21, 393–404, doi:10.1081/cbi-120038607.

35. Govic, A.; Levay, E.A.; Hazi, A.; Penman, J.; Kent, S.; Paolini, A.G. Alterations in male sexual behaviour, attractiveness and testosterone levels induced by an adult-onset calorie restriction regimen. Behav Brain Res 2008, 190, 140–146, doi:10.1016/j.bbr.2008.02.013.

36. Hull, E.M.; Dominguez, J.M. Sexual behavior in male rodents. Horm Behav 2007, 52, 45–55, doi:10.1016/j.yhbeh.2007.03.030.

37. Harding, S.M.; Velotta, J.P. Comparing the relative amount of testosterone required to restore sexual arousal, motivation, and performance in male rats. Horm Behav 2011, 59, 666–673, doi:10.1016/j.yhbeh.2010.09.009.

38. Pfaus, J.G. Pathways of sexual desire. J Sex Med 2009, 6, 1506–1533, doi:10.1111/j.1743-6109.2009.01309.x.

39. Xie, K.; Wang, C.; Scifo, E.; Pearson, B.; Ryan, D.; Henzel, K.; Markert, A.; Schaaf, K.; Mi, X.; Tian, X.;, et al. Intermittent fasting boosts sexual behavior by limiting the central availability of tryptophan and serotonin. Cell Metab 2025, 37, 1189–1205 e1187, doi:10.1016/j.cmet.2025.03.001.

40. Marino, M.D.; Cronise, K.; Lugo, J.N., Jr.; Kelly, S.J. Ultrasonic vocalizations and maternal-infant interactions in a rat model of fetal alcohol syndrome. Dev Psychobiol 2002, 41, 341–351, doi:10.1002/dev.10077.

41. Capone, F.; Bonsignore, L.T.; Cirulli, F. Methods in the analysis of maternal behavior in the rodent. Curr Protoc Toxicol 2005, *Chapter 13*, Unit13 19, doi:10.1002/0471140856.tx1309s26.

42. Koek, W.; Sandoval, T.L.; Daws, L.C. Effects of the antidepressants desipramine and fluvoxamine on latency to immobility and duration of immobility in the forced swim test in adult male C57BL/6J mice. Behav Pharmacol 2018, 29, 453–456, doi:10.1097/FBP.0000000000000371.

43. de Kloet, E.R.; Molendijk, M.L. Coping with the Forced Swim Stressor: Towards Understanding an Adaptive Mechanism. Neural Plast 2016, 2016, 6503162, doi:10.1155/2016/6503162.

44. Cora, M.C.; Kooistra, L.; Travlos, G. Vaginal Cytology of the Laboratory Rat and Mouse: Review and Criteria for the Staging of the Estrous Cycle Using Stained Vaginal Smears. Toxicol Pathol 2015, 43, 776–793, doi:10.1177/0192623315570339.

45. Koolhaas, J.M.; Coppens, C.M.; de Boer, S.F.; Buwalda, B.; Meerlo, P.; Timmermans, P.J. The resident-intruder paradigm: a standardized test for aggression, violence and social stress. J Vis Exp 2013, e4367, doi:10.3791/4367.

46. Lucio, R.A., Fuentes-Morales, M.R., Fernández-Guasti, A.. Copulation in Rats: Analysis of Behavioral and Seminal Parameters. In Animal Models of Reproductive Behavior. Neuromethods, Paredes, R.G., Portillo, W., Bedos, M, Ed.; Humana: New York, 2023; Volume 200.

47. Karabaşoğlu, C.; Erbaş, O. Rat Sexual Behaviours. The Journal of Experimental and Basic Medical Sciences 2021, 2, 139–146, doi:10.5606/jebms.2021.75650.

48. Blanchard, R.J.; Blanchard, D.C. Attack and defense in rodents as ethoexperimental models for the study of emotion. Prog Neuropsychopharmacol Biol Psychiatry 1989, 13 *Suppl*, S3–14, doi:10.1016/0278-5846(89)90105-x.

49. Kruschke, J.K. Bayesian analysis reporting guidelines. *Nat*. Hum. Behav. 2021, 5, 1282–1291, doi:10.1038/s41562-021-01177-7.

50. Vehtari, A.; Gelman, A.; Gabry, J. Practical Bayesian model evaluation using leave-one-out cross-validation and WAIC. Statistics and Computing 2017, 27, 1413–1432, 10.1007/s11222-016-9696-4.

51. Vehtari, A.; Gelman, A.; Simpson, D.; Carpenter, B.; Bürkner, P.-C. Rank-normalization, folding, and localization: An improved R^ for assessing convergence of MCMC (with discussion). Bayesian Anal. 2021, 16, 667–718, doi:10.1214/20-BA1221.

52. Gelman, A., Carlin, J.B., Stern, H.S., & Rubin, D.B.. Bayesian Data Analysis; Chapman and Hall/CRC: 1995.

53. Schwaferts, P.; Augustin, T. Bayesian decisions using regions of practical equivalence (ROPE): Foundations; University of Munich: 2020.

54. Kruschke, J.K. Rejecting or Accepting Parameter Values in Bayesian Estimation. Advances in Methods and Practices in Psychological Science 2018, 1, 270–280, doi:10.1177/2515245918771304.

55. Taylor-LaPole, A.M.; Cunny, H.C.; Shockley, K.R. Statistical Analysis of Rodent Body Weight Data is Robust to Departures from Normality in Historical National Toxicology Program Studies Dated 1980-2013. Journal of Young Investigators 2022, 25, 1–12.

