## Supplementary Materials for "Maternal preconception calorie restriction reprograms coping strategies, socio-sexual behaviour, and endocrine function in adult rat offspring"

### Supplementary 1 Male Female Interaction Clusters

| Aggressive | Dominance | Male Copulatory |
| --- | --- | --- |
| - Attack - Bite - Lateral Threat - Clinch Attack - Kicking - Standing Upright - Boxing/Nose Off | - Pinning/Keep Down - Walkover | - Mount - Intromission - Ejaculation |
| Female Copulatory | Social | Sociosexual |
| - Soliciting - Darting - Paracopulatory - Hopping - Ear wiggling - Consummatory - Rejection - Posturing - Nose-off | - Sniffing - Allogrooming | - Anogenital Sniffing - Pursuit/Follow/Chase |
| Non-social | Submissive |  |
| - Self-grooming - Rearing - Resting | - Flee |  |

### Supplementary 2 Model Specification Table

| Measurement/Test | Sample Size^1^ | Model Family*^2^* | Predictors*^3^* | Random Effects | ROPE Bound Decision*^4^* |
| --- | --- | --- | --- | --- | --- |
| Body Weight: F0 Females (Full Model) | N = 48  (48 females) | GAMM | Diet, Time, Phase | Individual | 10% body weight |
| Body Weight: F0 Females (Diet Phase Only Model) | N = 48  (48 females) | GLM | Diet, Time | Individual | Slope comparison to Control |
| Body Weight: Offspring | N = 368  (184 females,  184 males) | GLMM | Diet, PND | Individual | 10% body weight |
| Nest Construction/Quality | N = 45  (45 females) | Cumulative Logit Mixed-Effects Model (Ordinal) | Diet, Time, Time of Day (covariate) | Individual |  |
| Maternal Behaviour Observations | N = 45  (45 females) | GLM | Diet |  |  |
| Pup Retrieval Test | N = 45  (45 females) | GLM | Diet |  |  |
| Pup Developmental Milestones | N = 47  (47 females) | GLM | Diet |  |  |
| Locomotor Activity | N = 92  (46 females,  46 males) | GLMM | Diet, Sex, Session | Individual |  |
| Running Wheel Activity | N = 92  (46 females,  46 males) | GLM | Diet, Sex |  |  |
| Elevated Plus Maze | N = 108  (54 females,  54 males) | GLM | Diet, Sex, Distance Travelled (covariate) |  |  |
| Open Field Test | N = 112  (56 females,  56 males) | GLM | Diet, Sex, Distance Travelled (covariate) |  |  |
| Back Test (Escapes & Vocalisations) | N = 109  (53 females,  56 males) | GLMM | Diet, Sex, Session | Individual |  |
| Forced Swim Test | N = 109  (53 females,  56 males) | GLM | Diet, Sex |  |  |
| Sucrose Preference | N = 55  (27 females,  28 males) | GLM | Diet |  |  |
| Male–Female Interaction (All Behaviour Categories) | N = 50  (25 females,  25 males) | Multinomial Regression | Diet, Session |  | ±18% ROPE bound |
| Male Copulatory Bias vs Aggression | N = 50  (25 females,  25 males) | Multinomial Regression | Diet, Session |  | ±18% ROPE bound |
| Resident–Intruder | N = 53  (53 males) | GLM | Diet |  |  |
| Basal Serum Testosterone | N = 56  (56 males) | GLM | Diet |  |  |
| Faecal Testosterone Metabolites | N = 41  (41 males) | GLM | Diet, Time | Individual |  |
| *^1^* Animals were excluded from specific analyses only where predefined technical or welfare-related criteria were met (e.g. Equipment malfunction, incomplete task engagement, or data acquisition failure). No animals were excluded based on treatment group or outcome  *^2^* GAMM = Generalised Additive Mixed Model; GLMM = Generalised Linear Mixed Model; GLM = Generalised Linear Model | | | | | |
| *^3^* Predictors include all interactions unless otherwise specified | | | | | |
| *^4^* Blank ROPE Bound Decision indicates standard ROPE formulation was used (e.g. 0.1 x standard deviation of sample) | | | | | |

### Supplementary 3 Model Specification, Regression Summary and Diagnostics Results Online Repository Link

Link to online repository:

<https://www.epigenes.com.au/our-research/maternal-preconception-dietary-restriction>

### Supplementary 4 Count per Session for Social Behavioural Clusters

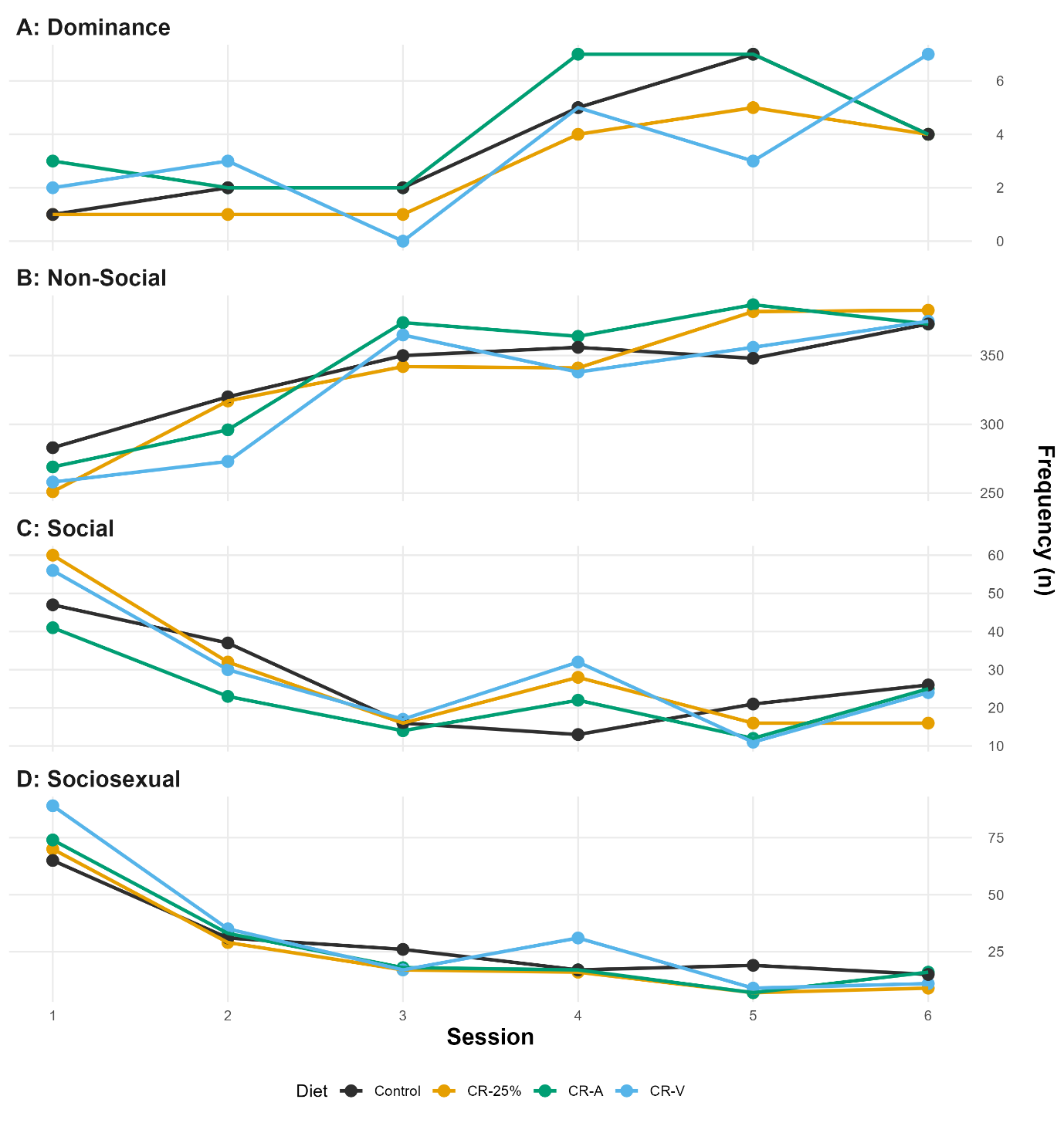

Figure 1 Frequencies of male–female social behaviours across days by treatment group. Line plots show the observed frequency of behaviours from four functional clusters: Aggressive, Female Rejection, Male Copulatory, and Submissive, recorded over six sessions. Each panel corresponds to one behavioural cluster, with lines representing treatment groups (Control, CR-25%, CR-A, CR-V). Variability in patterns is apparent across both behaviours and groups, particularly in copulatory interactions, where peak frequencies occurred early in the observation period.

### Supplementary 5 Male Female Interaction ROPE Contrasts by Day: Dominance Behaviours

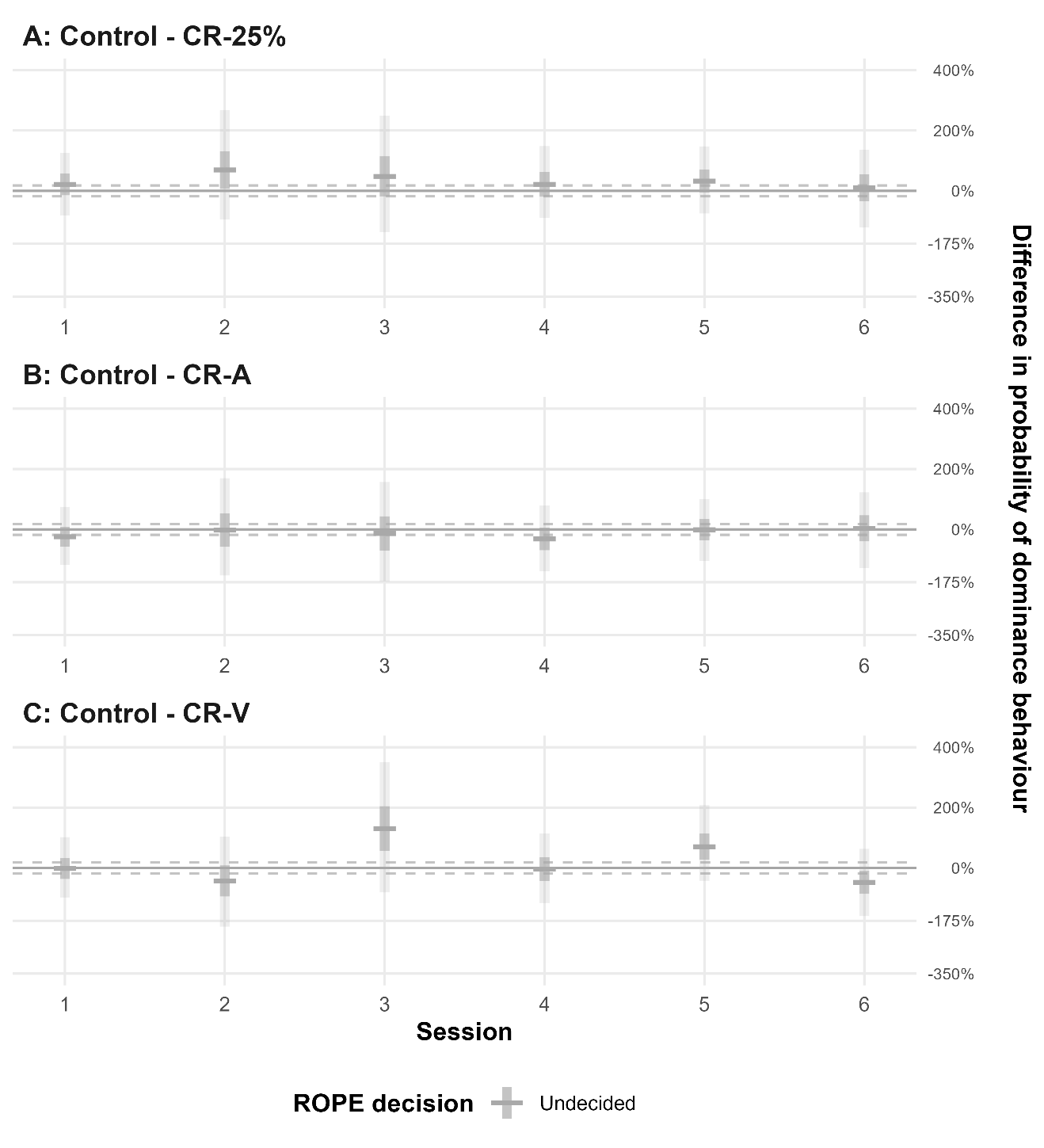

Figure 2 ROPE analysis contrast plots over time for differences in dominance behaviour. This figure displays the estimated differences in the probability of dominance behaviour between pairs of treatment groups at six time points (Sessions 1-6). Horizontal lines indicate posterior means, with darker vertical shaded areas representing 50% credible intervals and lighter areas representing 95% credible intervals. ROPE range: 0 ±18%. Positive values indicate a higher probability of dominance in the first group of the contrast compared to the second.

### Supplementary 6 Male Female Interaction ROPE Contrasts by Day: Female Copulatory Behaviours

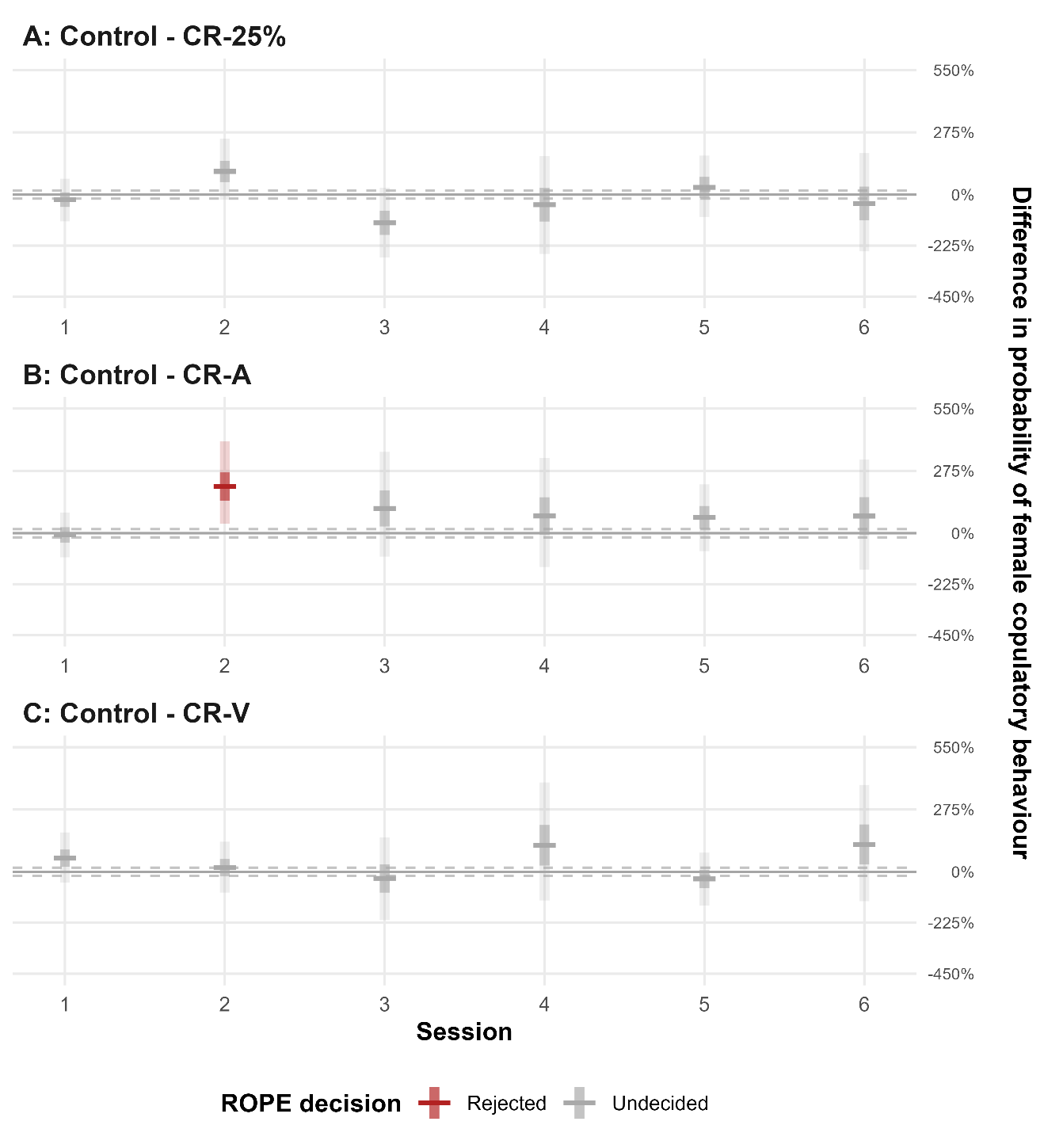

Figure 3 ROPE analysis contrast plots over time for differences in female copulatory behaviour. This figure displays the estimated differences in the probability of female copulatory behaviour between pairs of treatment groups at six time points (Sessions 1-6). Horizontal lines indicate posterior means, with darker vertical shaded areas representing 50% credible intervals and lighter areas representing 95% credible intervals. ROPE range: 0 ±18%. Positive values indicate a higher probability of female copulatory behaviour in the first group of the contrast compared to the second.

### Supplementary 7 Male Female Interaction ROPE Contrasts by Day: Social Behaviours

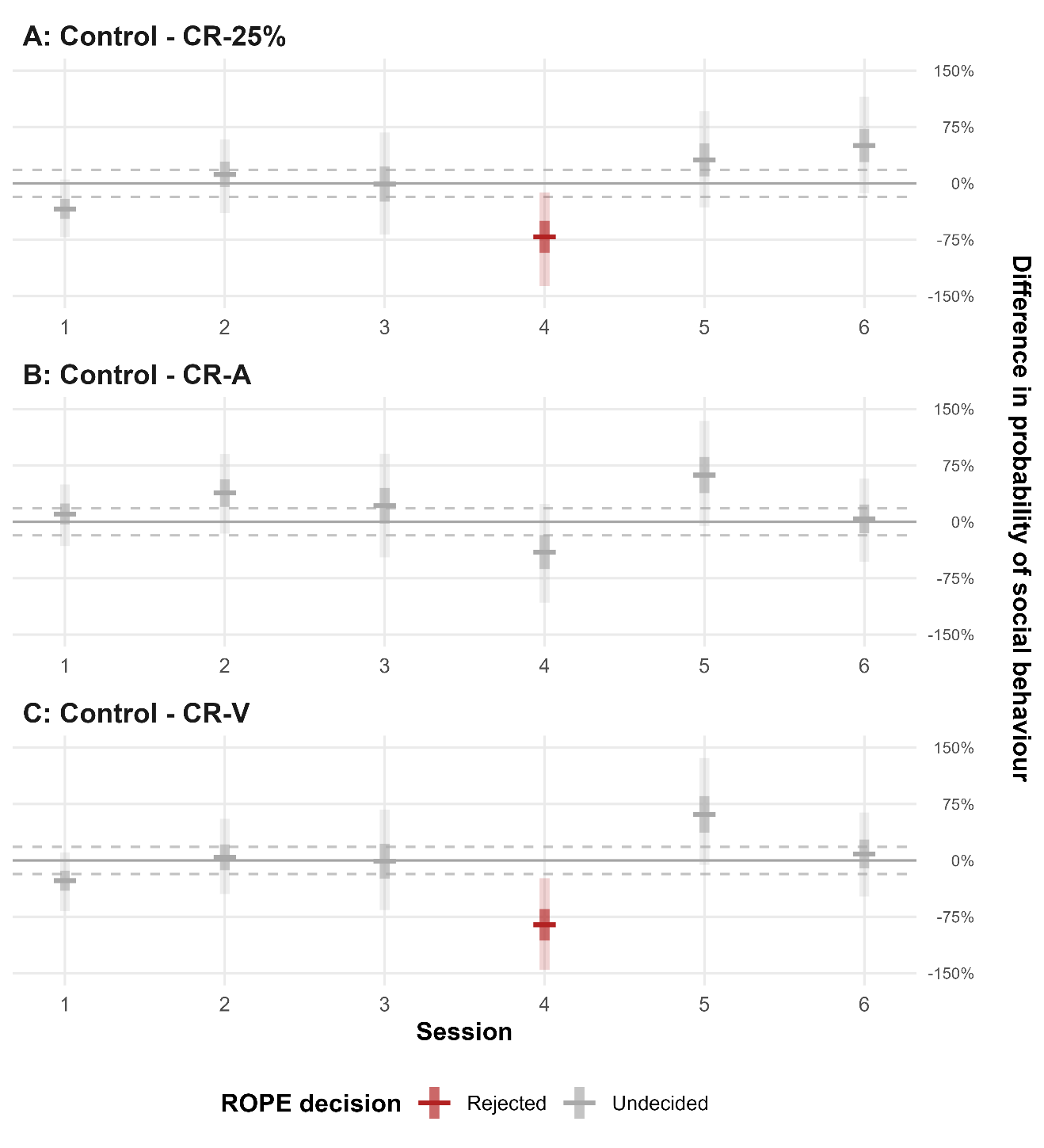

Figure 4 ROPE analysis contrast plots over time for differences in social behaviour. This figure displays the estimated differences in the probability of social behaviour between pairs of treatment groups at six time points (Sessions 1-6). Horizontal lines indicate posterior means, with darker vertical shaded areas representing 50% credible intervals and lighter areas representing 95% credible intervals. ROPE range: 0 ±18%. Positive values indicate a higher probability of social in the first group of the contrast compared to the second.

### Supplementary 8 Male Female Interaction ROPE Contrasts by Day: Sociosexual Behaviours

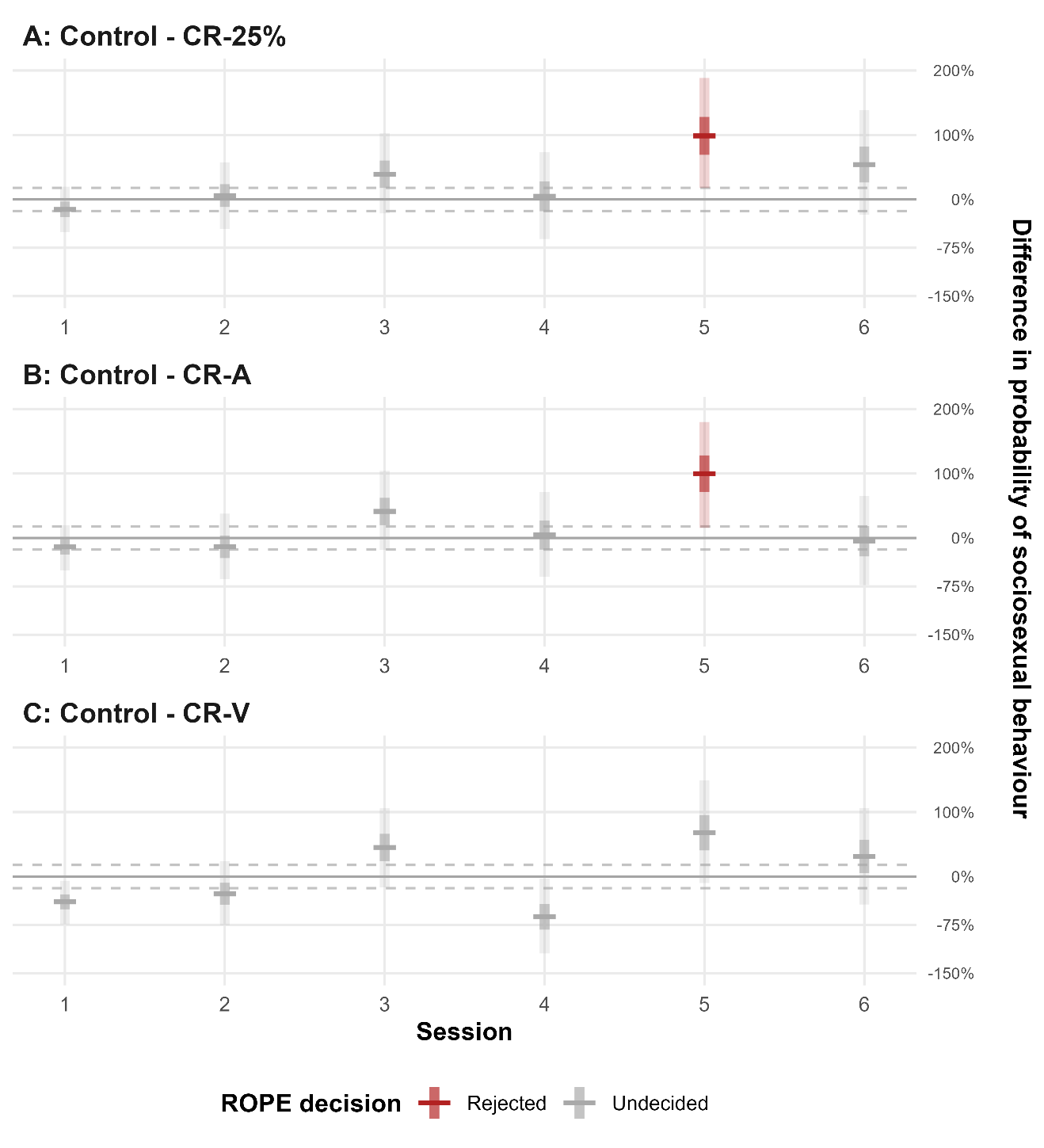

Figure 5 ROPE analysis contrast plots over time for differences in sociosexual behaviour. This figure displays the estimated differences in the probability of sociosexual behaviour between pairs of treatment groups at six time points (Sessions 1-6). Horizontal lines indicate posterior means, with darker vertical shaded areas representing 50% credible intervals and lighter areas representing 95% credible intervals. ROPE range: 0 ±18%. Positive values indicate a higher probability of sociosexual in the first group of the contrast compared to the second.

### Supplementary 9 Male Female Interaction ROPE Contrasts by Day: Submissive Behaviours

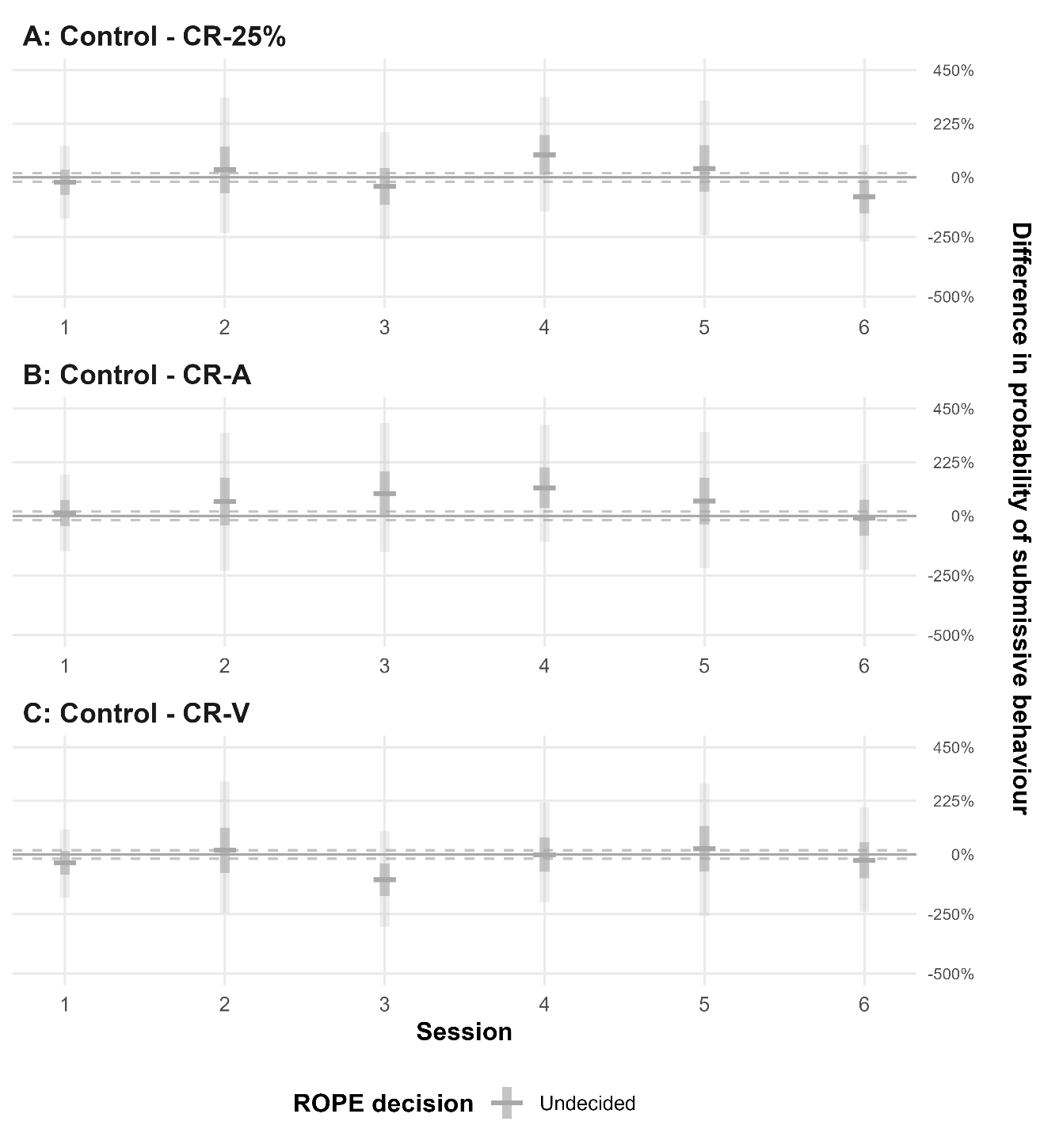

Figure 6 ROPE analysis contrast plots over time for differences in submissive behaviour. This figure displays the estimated differences in the probability of submissive behaviour between pairs of treatment groups at six time points (Sessions 1-6). Horizontal lines indicate posterior means, with darker vertical shaded areas representing 50% credible intervals and lighter areas representing 95% credible intervals. ROPE range: 0 ±18%. Positive values indicate a higher probability of submissive in the first group of the contrast compared to the second.

### Supplementary 10 Male Female Interaction ROPE Contrasts by Day: Nonsocial Behaviours

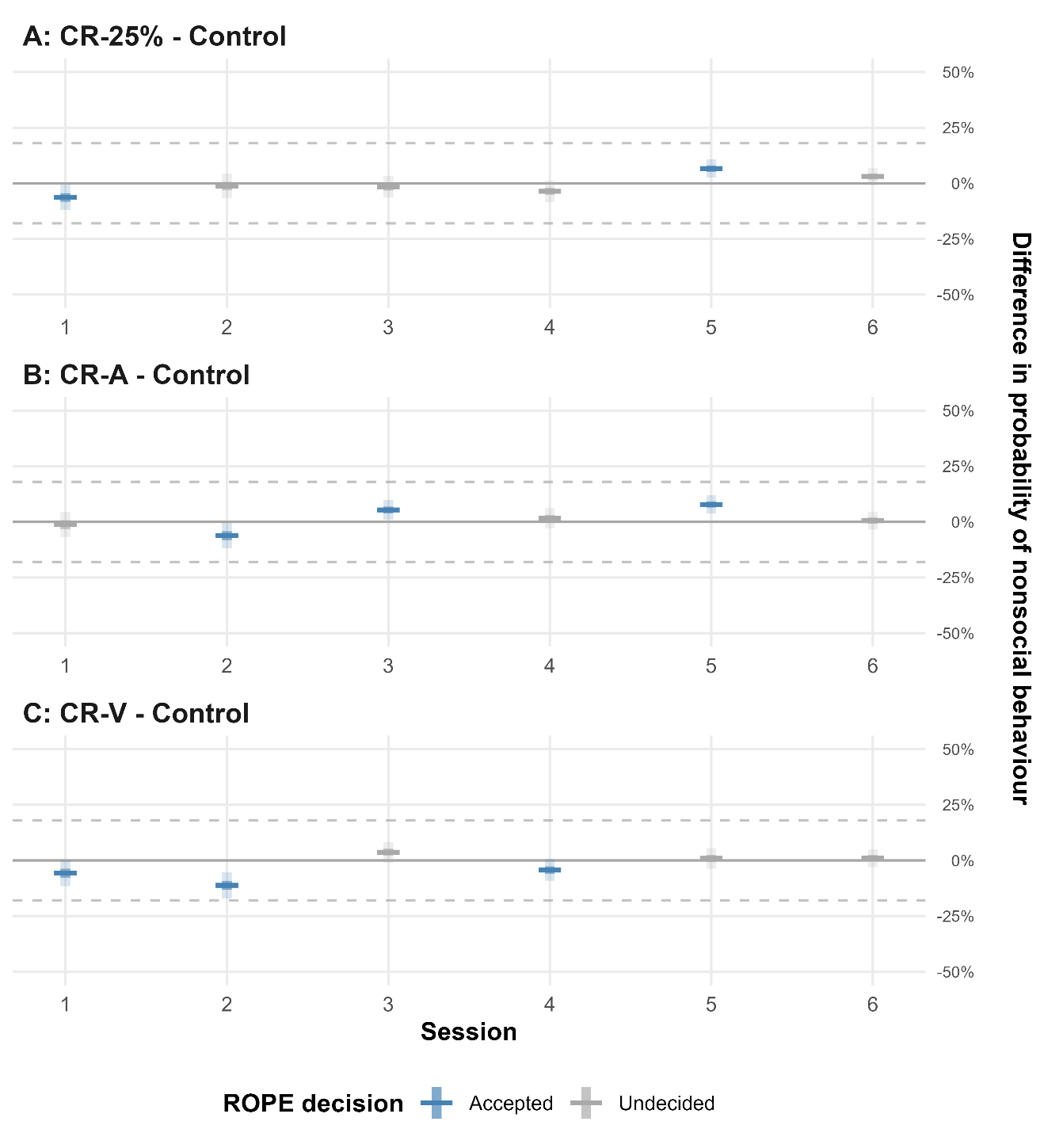

Figure 7 ROPE analysis contrast plots over time for differences in nonsocial behaviour. This figure displays the estimated differences in the probability of nonsocial behaviour between pairs of treatment groups at six time points (Sessions 1-6). Horizontal lines indicate posterior means, with darker vertical shaded areas representing 50% credible intervals and lighter areas representing 95% credible intervals. ROPE range: 0 ±18%. Positive values indicate a higher probability of nonsocial in the first group of the contrast compared to the second.
